# Perturbations of transcription and gene expression-associated processes alter distribution of cell size values in *Saccharomyces cerevisiae*

**DOI:** 10.1101/461210

**Authors:** Nairita Maitra, Jayamani Anandhakumar, Heidi M. Blank, Craig D. Kaplan, Michael Polymenis

**Affiliations:** Department of Biochemistry and Biophysics, Texas A&M University, College Station, TX 77843

**Keywords:** cell size/, gamma distribution/, RSC/, RNA polymerase/, THO

## Abstract

The question of what determines whether cells are big or small has been the focus of many studies because it is thought that such determinants underpin the coupling of cell growth with cell division. In contrast, what determines the overall pattern of how cell size is distributed within a population of wild type or mutant cells has received little attention. Knowing how cell size varies around a characteristic pattern could shed light on the processes that generate such a pattern and provide a criterion to identify its genetic basis. Here, we show that cell size values of wild type *Saccharomyces cerevisiae* cells fit a gamma distribution, in haploid and diploid cells, and under different growth conditions. To identify genes that influence this pattern, we analyzed the cell size distributions of all single-gene deletion strains in *Saccharomyces cerevisiae.* We found that yeast strains which deviate the most from the gamma distribution are enriched for those lacking gene products functioning in gene expression, especially those in transcription or transcription-linked processes. We also show that cell size is increased in mutants carrying altered activity substitutions in Rpo21p/Rpb1, the largest subunit of RNA polymerase II (Pol II). Lastly, the size distribution of cells carrying extreme altered activity Pol II substitutions deviated from the expected gamma distribution. Our results are consistent with the idea that genetic defects in widely acting transcription factors or Pol II itself compromise both cell size homeostasis and how the size of individual cells is distributed in a population.

## INTRODUCTION

Mechanisms that control cell size have long been viewed as critical for the coupling between cell growth and cell division, which in turn governs rates of cell proliferation (Turner et al. 2012; Pringle 1981; Ginzberg et al. 2015; Westfall and Levin 2017; Willis and Huang 2017). Hence, size control has attracted attention in many systems, from bacteria and yeasts to animals (Si et al. 2017; Tzur et al. 2009; Son et al. 2012; Jorgensen et al. 2002; Zhang et al. 2002). Most studies have dealt with situations where the typical size of cells in a given experimental system and condition shifts to a different value, due to genetic or environmental perturbations. Despite many rounds of cell division, proliferating cells usually maintain their size in a given nutrient environment. Considering cell size as a proxy for cell growth, then shifts to a smaller or larger size provide a convenient 'metric' to gauge alterations in biological processes that are thought to be central to the physiological coupling between cell growth and division.

Cells tune their gene expression output to their size, to maintain the proper concentrations of macromolecules as cells change in volume (Vargas-Garcia et al. 2018). Changes in ploidy and the well-known positive association between cell size and DNA content (Gregory 2001) perhaps illustrate a straightforward solution to this problem. Compared to smaller haploid and diploid cells, larger polyploid ones have more genomic templates from which to drive gene expression. It has also been reported that ploidy-associated increases in cell size drive transcriptional changes (Galitski et al. 1999; Wu et al. 2010). The situation appears more complex in cells of different size but of the same genome (Marguerat and Bahler 2012; Zhurinsky et al. 2010). In fission yeast, it has been proposed that cells of different size regulate global transcription rates regardless of cellular DNA content so that their transcriptional output per protein remains constant (Zhurinsky et al. 2010). Based on single molecule transcript counting in mammalian cells, a positive association between transcription burst magnitude and cell size has been reported (Padovan-Merhar et al. 2015). Furthermore, it appears that the doubling of the available DNA templates for transcription after DNA replication is countered by a decrease in transcription burst frequency in cells that have replicated their DNA, later in the cell cycle (Padovan-Merhar et al. 2015). These mechanisms, involving independent control of the frequency and the magnitude of transcription bursts, are thought to maintain the scaling of mRNA counts with the size of mammalian cells. In the budding yeast *Saccharomyces cerevisiae,* analogous single molecule experiments monitoring transcription bursts as a function of cell size and cell division have not been reported. Instead, a somewhat different mechanism has been proposed to explain the positive association of mRNA steady-state levels with cell size, due to increased stability of mRNAs in larger cells (Mena et al. 2017). Furthermore, it has been proposed that levels of active RNA Pol II are higher in small G1 cells with un-replicated DNA (Mena et al. 2017).

The genetic control of cell size has been studied extensively in *S. cerevisiae.* In this organism, systematic surveys of all single-gene deletions have been carried out to identify mutants that are bigger or smaller than the wild type (Zhang et al. 2002; Jorgensen et al. 2002). Similar size-based screens have also been carried out in other organisms (Bjorklund et al. 2006), with similar outputs, namely the identification of small or large-celled mutants. In contrast, much less attention has been placed on how the size of individual cells within a population, mutant or wild type, is distributed. We reasoned that if we might first determine if yeast cells fit a particular distribution of sizes, we might then determine what types of mutants alter such a stereotypical distribution of cell sizes to understand its genetic basis.

Here we report that size in a population of *S. cerevisiae* cells is best described with a gamma distribution. We also identify genes that are required to maintain this distribution. These genes overwhelmingly encode proteins involved in global gene expression, especially in transcription. Lastly, we show that defects arising from alterations to the Pol II active site alter size homeostasis and the pattern of size distributions.

## MATERIALS AND METHODS

### Yeast strains and cell size measurements

For cell size measurements we report in Figure 5, the homozygous deletion strains were in the diploid S288C background of strain BY4743 *(MATα/α hiS4Δ1/hiS4Δ1 leu2Δ0/leu2Δ0 LYS2/lys2Δ0 met15Δ0/MET15 urα3Δ0/urα3Δ0).* The strains were constructed by the Systematic Deletion Project (Giaever et al. 2002; Brachmann et al. 1998).

For cell size measurements we report in Figure 6, genomic variants of *RPO21/RPB1* were created by CRISPR/Cas9-mediated genome editing (see Table S1 for a list of the oligonucleotides used in this study) of strain CKY3284, a derivative of the S288c strain background (see Table S2 for a list of the strains used in this study). Briefly, the sequence encoding a guide RNA was cloned into plasmid pML107 ((Laughery et al. 2015); see Table S3 for a list of the plasmids used in this study). This plasmid was introduced into CKY3284 by transformation, along with annealed double-stranded repair oligonucleotides or annealed overlapping oligonucleotides (Integrated DNA Technologies, Skokie, Illinois; see Table S1) filled in with Phusion DNA polymerase (New England Biolabs, Ipswich, Massachusetts) containing either a silently mutated PAM site or both a silently mutated PAM site and relevant *RPO21/RPB1* mutation. Variants were confirmed by PCR amplification of the mutated region and DNA sequencing. Oligonucleotides were annealed as follows: synthesized, lyophilized oligonucleotides were resuspended at 100 µM in 10 mM Tris pH 8.0. Perfectly matched oligonucleotide pairs were annealed as follows: 37.5 µl of each oligonucleotide was mixed with 25 µl of 1M Tris pH 8.0 and heated for 5 minutes at 95°, then tubes were transferred into a 70° heat block followed by removal of heat block from heater; they were moved to 4° overnight when block temperature reached room temperature. 8 µl of these annealed oligonucleotides were used as repair templates in individual transformations. Oligonucleotides that were overlapping (noted in Table S3) were annealed and extended by 5 cycles of standard PCR followed by thermal annealing as above.

All strains were cultured at 30° in standard YPD medium (1% ^w^/v yeast extract, 2% ^w^/v peptone, 2% ^w^/v dextrose). Cell size was measured with a Z2 Beckman Coulter channelyzer as described previously (Guo et al. 2004; Bogomolnaya et al. 2006).

DNA content analysis was done as we have described previously (Hoose et al. 2012; Hoose et al. 2013).

### Statistical analysis

In all our statistical analyses we used R language packages, as indicated in each case. The cell size frequency distributions we analyzed from the literature were from (Soma et al. 2014) for the BY4743 dataset, and from (Jorgensen et al. 2002) for the haploid dataset (BY4741; *MATα hiS4Δ1 leu2Δ0 met15Δ0 urα3Δ0)* (see File S1; sheets 'by4743_raw', and 'Jorgensen_raw', respectively). Replicates of several strains in the Jorgensen dataset were marked as such, following their systematic ORF name. The cell size frequencies from (Soma et al. 2014) and (Jorgensen et al. 2002) were used to simulate distributions from n=1000 cells in every case (see File S1; sheets 'by4743', and 'jorgensen', respectively). Similarly, we also generated the size distributions shown in Figures 5 and 6 ((see File S1; sheets 'figure5', and 'figure6', respectively). To generate counts from frequencies for downstream statistical analysis, we used the R code listed in File S2.

To test for normality, we implemented the Shapiro-Wilk test (Shapiro and Wilk 1965) from the *stats* R language package, as described in detail in File S2. The corresponding p-values are shown in File S1, in the sheet columns marked as 'SW(p)'. Since normality was not observed for any of the BY4743-based mutant distributions, we then fitted them to several right-side skewed distributions, including lognormal, Gamma and Weibull. To this end, we used the goodness-of-fit function 'gofstat' of the *fitdistrplus* R language package (Delignette-Muller and Dutang 2015), implementing the Anderson-Darling test (Anderson and Darling 1952) for each of the samples shown in Table 1. Using the goodness-of-fit function 'gofstat' of the fitdistrplus R language package we also obtained the corresponding statistic values for the Kolmogorov-Smirnov and Cramér-von Mises tests (Table 1). Fitting wild type cell size distributions to more complex, three-parameter generalized gamma models only minimally improved the fit, but it increased complexity. As a result, the preferred model was the standard two-parameter gamma distribution, based on a lower value of the Bayesian Information Criterion (Schwarz 1978). To calculate the shape (α) and rate (β) parameters of the gamma-fitted distributions (see Table 1), we used the maximum-likelihood estimates approach implemented by the 'fitdistr' function of the *MASS* R language package. The same analysis was applied to the two BY4741 samples from the 'jorgensen' dataset shown in Table 1. To obtain the Anderson-Darling test p-values for gamma distribution fits for each strain, we used the 'gofTest' function of the *goft* R language package (González-Estrada and Villaseñor 2018). These p-values are shown in File S1, in the sheet columns marked as 'AD(p)'. For the 'jorgensen' dataset we also used the 'ad.test' function of the *goftest* R language package, as follows: ad.test(strain, null = “dgamma”, shape = 3.8277, rate = 0.078949). The shape and rate parameters were the average of the two wild type BY4741 samples in the 'jorgensen' dataset. We also used the same functions to obtain the test's statistic, which was used to identify the 49 genes that when deleted yield size distributions that deviate the most from a gamma pattern (shown in File S1/sheet 'Gamma_deviant_Genes').

**Table 1.** Statistical parameters of cell size distributions of wild type strains

| Strain | Carbon source | Anderson-Darling Statistic |  |  | Gamma parameters |  |
| --- | --- | --- | --- | --- | --- | --- |
| | | Lognormal | Weibull | Gamma | Shape ($\alpha$) | Rate ($\beta$) |
| BY4743 | Dextrose(2%) | 3.161116 | 2.479761 | 0.769909 | 6.624628 | 0.060432 |
| BY4743 | Dextrose(2%) | 2.481744 | 3.453834 | 0.907658 | 6.839326 | 0.0613 |
| BY4743 | Dextrose(2%) | 2.350162 | 4.639409 | 1.397938 | 7.939692 | 0.062422 |
| BY4743 | Dextrose(2%) | 9.379353 | 3.256118 | 2.080058 | 6.327387 | 0.051845 |
| BY4743 | Dextrose(0.5%) | 2.400366 | 5.534772 | 1.885079 | 6.631854 | 0.056427 |
| BY4743 | Dextrose(0.5%) | 2.681816 | 4.711678 | 1.737208 | 7.028829 | 0.059809 |
| BY4743 | Dextrose(0.5%) | 2.554072 | 4.049485 | 1.352729 | 6.523879 | 0.056879 |
| BY4743 | Dextrose(0.5%) | 2.238338 | 4.465921 | 1.296817 | 6.682971 | 0.05811 |
| BY4743 | Dextrose(0.5%) | 1.657507 | 5.112165 | 1.088213 | 6.92371 | 0.059789 |
| BY4743 | Dextrose(0.5%) | 2.257491 | 4.7403 | 1.40862 | 7.118115 | 0.060818 |
| BY4743 | Dextrose(0.1%) | 2.093822 | 3.809769 | 0.68981 | 5.491663 | 0.051904 |
| BY4743 | Dextrose(0.1%) | 3.495197 | 2.657985 | 1.090478 | 5.502986 | 0.050511 |
| BY4743 | Dextrose(0.1%) | 0.998583 | 5.800609 | 0.817631 | 5.038751 | 0.050269 |
| BY4743 | Dextrose(0.1%) | 1.429023 | 4.734133 | 0.725159 | 4.987775 | 0.049489 |
| BY4743 | Dextrose(0.05%) | 3.090379 | 4.551848 | 0.58666 | 4.084743 | 0.043347 |
| BY4743 | Dextrose(0.05%) | 1.194635 | 5.519737 | 0.903827 | 4.601468 | 0.047501 |
| BY4743 | Dextrose(0.05%) | 2.240336 | 5.114651 | 0.608227 | 4.13688 | 0.046411 |
| BY4743 | Dextrose(0.05%) | 1.685737 | 5.903186 | 0.693725 | 4.137698 | 0.046938 |
| BY4743 | Dextrose(0.05%) | 2.300439 | 3.695214 | 0.388641 | 4.936219 | 0.052235 |
| BY4743 | Dextrose(0.05%) | 1.850587 | 4.21797 | 0.692334 | 4.86957 | 0.04992 |
| BY4743 | Galactose(2%) | 4.48614 | 2.026274 | 1.063566 | 4.352128 | 0.049247 |
| BY4743 | Galactose(2%) | 4.019618 | 1.957646 | 0.849565 | 4.34835 | 0.048053 |
| BY4743 | Galactose(2%) | 2.866003 | 3.179497 | 0.85723 | 4.460477 | 0.050979 |
| BY4743 | Galactose(2%) | 3.541828 | 2.324092 | 0.906612 | 4.651173 | 0.053049 |
| BY4743 | Galactose(2%) | 5.23192 | 1.70193 | 1.398188 | 4.561268 | 0.0503 |
| BY4743 | Galactose(2%) | 4.898133 | 2.389382 | 1.858014 | 4.886206 | 0.053239 |
| BY4743 | Glycerol(2%) | 4.023934 | 1.706263 | 0.484643 | 4.576071 | 0.046215 |
| BY4743 | Glycerol(2%) | 4.912524 | 1.426551 | 0.417717 | 4.19207 | 0.046484 |
| BY4743 | Glycerol(2%) | 3.537731 | 2.212626 | 0.166045 | 4.243508 | 0.04657 |
| BY4741 | Dextrose(2%) | 8.884836 | 1.728113 | 1.143929 | 3.651767 | 0.074363 |
| BY4741 | Dextrose(2%) | 9.115701 | 3.162732 | 1.503567 | 4.003634 | 0.083534 |

All other R language functions and packages used to generate plots are described in the corresponding figures.

### Data Availability

Table S1 lists the oligonucleotides used in this study. Strains (Table S2) and plasmids (Table S3) are available upon request. File S1 contains all the datasets, including distributions of size frequencies and associated p-values, used in this study. The code used to analyze the data is provided in File S2. We have uploaded all the supplementary material (which includes seven supplementary figures) to figshare (https://figshare.com/s/999db4b2cb8c4deedbbcd).

## RESULTS

### Rationale

The approach we followed in this study is shown schematically in Figure 1. How different values of a single parameter are distributed can be instructive for the underlying processes leading to its distribution pattern. Accurately fitting the measured variable to a univariate distribution is also necessary for its proper statistical analysis, when determining how removed a given observation *(e.g.,* a mutant) is from the most typical one (wild type) in a population. With regards to the size of individual organisms, the usual pattern is that deviations from the common type are not symmetrical. Instead, small individuals tend to be more frequent than large ones, leading to distributions which are positively skewed, with a right-side tail (Frank 2016). The sizes of bacterial and animal cells have been modeled on lognormal distributions (Hosoda et al. 2011). To our knowledge, although *S. cerevisiae* is a prime model system in studies of size control, how size is distributed in this organism has not been examined. Consequently, our objectives for this study were: First, determine how cell size is distributed in *S. cerevisiae* (Figure 2). Second, use the distribution model that fits the best to empirical data as a metric to identify mutants that deviate the most from that distribution (Figures 3, 4). Third, validate the outliers experimentally (Figure 5), and test the role of the corresponding biological processes in determining size distributions (Figure 6).

**FIGURE 1.**
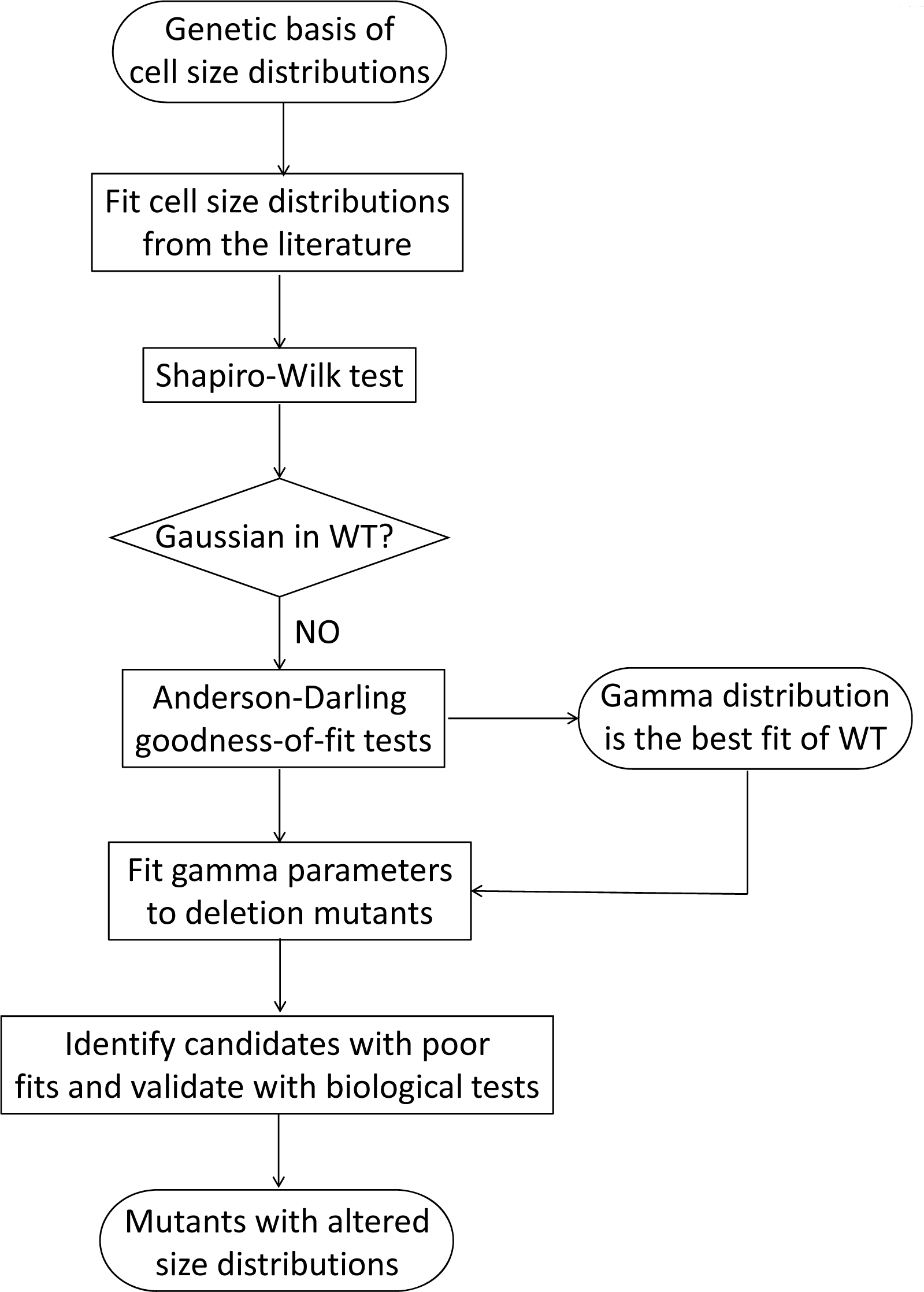
Flowchart of our approach to identify *S. cerevisiae* mutants with altered cell size distributions. See text for details.

### Cell size in *S. cerevisiae* fits a gamma distribution

We examined size frequency distributions from diploid cells cultured in different carbon sources, using published data from our laboratory (Soma et al. 2014). First, we looked at whether these distributions fit a Gaussian pattern. To test for normality, we employed the Shapiro-Wilk test (Shapiro and Wilk 1965), because it has the highest power compared to other tests (Razali and Wah 2011). In every one of the 29 wild type distributions from diploid cells we tested, the associated p-value was significantly lower than an alpha level of 0.01 (see Figure 2A, left box; the individual values are shown in File S1/sheet 'by4743_SW_AD_p'/column 'SW(p)'). Hence, the null hypothesis that these populations were normally distributed was rejected.

**FIGURE 2.**
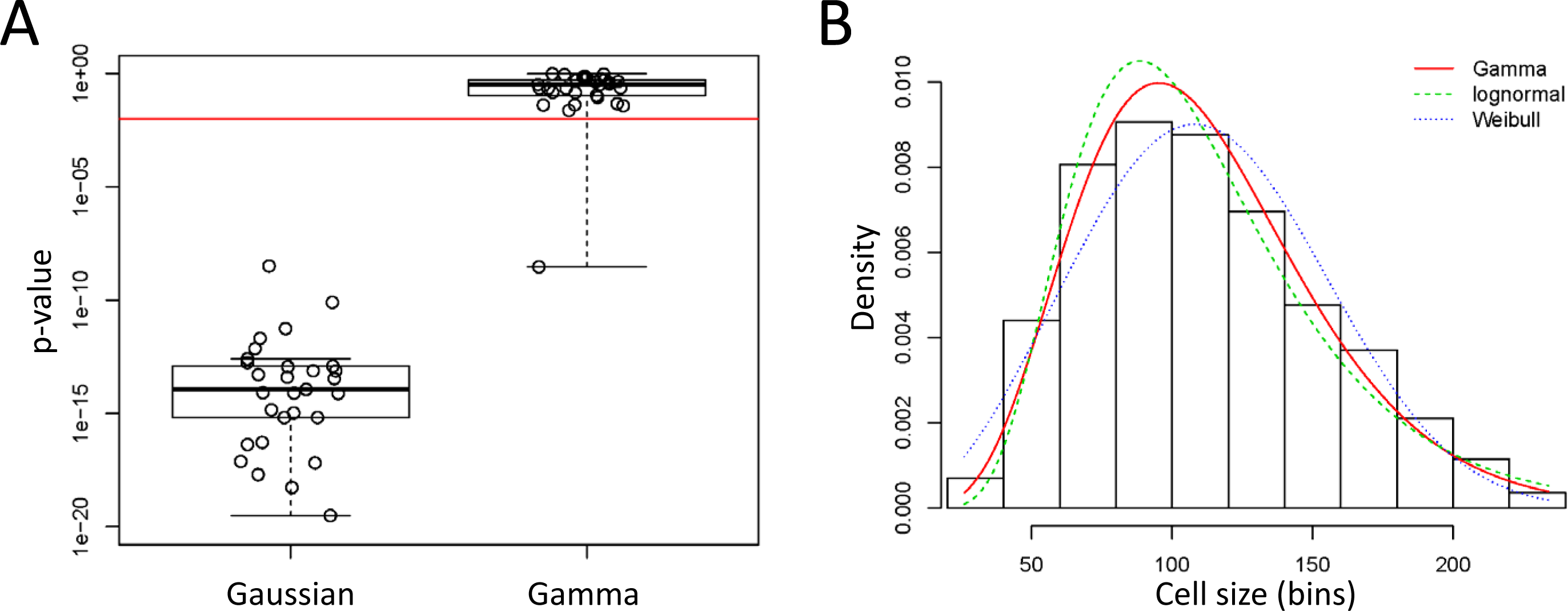
Cell size values of *S. cerevisiae* cells fit a Gamma distribution pattern. **A**, Box plots of the p-values associated with statistical tests of whether cell size values from (Soma et al. 2014) of wild type BY4743 cells (see Table 1) are distributed according to a normal (Gaussian), or Gamma distribution. The Shapiro-Wilk test was used to test for the Gaussian distribution, while the Anderson-Darling test was used to test for the Gamma distribution (see Materials and Methods). The red horizontal line indicates a significance level of p=0.01. The density plot of the sole outlier that did not fit a Gamma distribution is shown in Figure S1. **B**, Histogram and theoretical densities for the indicated cell size distribution of *S. cerevisiae* cells. The distributions were fitted to continuous, empirical data depicted in the histogram from wild type diploid (BY4743 strain) cells, cultured in standard YPD medium (1% yeast extract, 2% peptone, 2% dextrose). On the y-axis are the density frequency values, while on the x-axis are the cell size bins, encompassing the cell size values shown in the corresponding spreadsheet associated with this plot (see File S1). The plots were generated with the 'denscomp' function of the *fitdistrplus* R language package. Additional goodness-of-fit plots associated with this graph are shown in Figure S2.

Given that cell size distributions were positively skewed with right-side tails, we fit the empirical data to non-Gaussian distributions that yield such patterns, such as lognormal, Weibull and gamma (Table 1). To test the goodness of these fits, we primarily relied on the Anderson-Darling test (Anderson and Darling 1952), which is thought to be of higher power than other goodness-of-fit tests for non-normal distributions. In every case, the value of the test's statistic was the lowest for the gamma distribution (Table 1). To test further a lognormal distribution of these samples, we log-transformed these values and then examined if they were normally distributed, a prediction for values that are lognormally distributed. In no sample was this the case (not shown), arguing against lognormal distributions being the best fit for *S. cerevisiae* cell size values. Next, we calculated the Anderson-Darling associated p-value for a gamma distribution for each of the 29 diploid size distributions. In all but one sample, the p-value was higher than an alpha level of 0.01 (see Figure 2A, right box; the individual values are shown in File S1/sheet 'by4743_SW_AD_p'/column 'AD(p)'). Hence, the fits of all these samples are consistent with a gamma distribution. We looked at the empirical data of the one sample for which the p-value was significantly lower than the 0.01 cutoff (Figure S1). It appears that this distribution is irregular, with a shoulder on the right-side tail, perhaps explaining the poor fit (Figure S1). Nonetheless, even for this sample, the gamma distribution was the better fit, compared to lognormal or Weibull distributions (Table 1).

The suitability of a gamma distribution pattern to accurately describe *S. cerevisiae* cell size data was also evident when different (Gamma, lognormal, Weibull) theoretical fits were displayed on a histogram of continuous empirical data (Figure 2B; File S1, from the second sample in sheet 'by4743'). From the associated goodness-of-fit diagnostic plots (Figure S2), the gamma distribution is a better fit than the related lognormal and Weibull distributions for the values in the middle of the distribution (Figure S2C). For the data in the left-side tail, lognormal and gamma are superior to Weibull, albeit Weibull performs better for the data in the right-side tail of the distribution (Figure S2B). Taken together, the sizes of *S. cerevisiae* cells best fit a gamma distribution. In the Discussion, we expand on this finding.

Next, we calculated the shape (α) and rate (β) parameters that describe gamma distributions for each of the above wild type distributions (Table 1; see Materials and Methods). Note that the samples we analyzed were from cells growing in different carbon sources (dextrose, galactose, glycerol) and, in the case of dextrose, at different concentrations (0.05% to 2%) of this preferred carbon source for the organism. In all cases, the best fit of the size data was a gamma distribution (Table 1), regardless of nutrient composition. We note that the shape parameter (α) was reduced from 6.3-7.9 in rich, replete medium (2% dextrose) to 4.1-4.9 in carbon-restricted medium (0.05% dextrose; see Table 1), as expected for the accompanying reduction in cell size in this medium (Soma et al. 2014).

### Identifying mutants with cell size distributions that deviate the most from gamma

A significant outcome of our results that wild type cell size distributions from *S. cerevisiae* are best described with gamma distributions is that fitting an empirical distribution to a gamma pattern can be used as a 'metric' to identify mutants that deviate the most. To this end, we used the frequency data of cell distributions from (Jorgensen et al. 2002), which surveyed strains carrying single deletions in all non-essential genes in *S. cerevisiae* (Giaever et al. 2002). These strains were haploid, but in the same (S288c) genetic background as the strains we examined in Figure 2. They were also cultured in the same standard, dextrose-replete, YPD medium (see Materials and Methods). From the 5,052 size distributions in the (Jorgensen et al. 2002) dataset, not a single one fit to a normal, Gaussian distribution, based on the p-values from the Shapiro-Wilk test (Figure 3A, left box; the individual values are shown in File Sl/sheet 'jorgensen_SW_AD_p'/column 'SW(p)'). The best fits of the two wild type (BY4741) samples in the (Jorgensen et al. 2002) dataset were also gamma distributions, compared to lognormal or Weibull (Table 1; the bottom two rows). We note that since the size of both haploid and diploid cells fit a gamma distribution pattern (Table 1), ploidy per se does not appear to change the pattern of cell size distributions.

Next, we calculated the Anderson-Darling associated p-value for a gamma distribution for each of the 5,052 samples. For about half of them (n=2,527), the p-value was higher than an alpha level of 0.01 (see Figure 3A, right box; the individual values are shown in File Sl/sheet 'by4743_SW_AD_p'/column 'AD(p)'). Since even small experimental irregularities in empirical distributions disturb their fit to theoretical densities (e.g., see Figure S1), it is noteworthy that half of the frequencies could be adequately fitted. Furthermore, even for the samples whose p-values did not pass the alpha level of 0.01, it is clear for the overwhelming majority of them that their gamma-fitted p-values were orders of magnitude higher than their p-values for fits to Gaussian distributions (compare the right to the left plot in Figure 3A).

**FIGURE 3.**
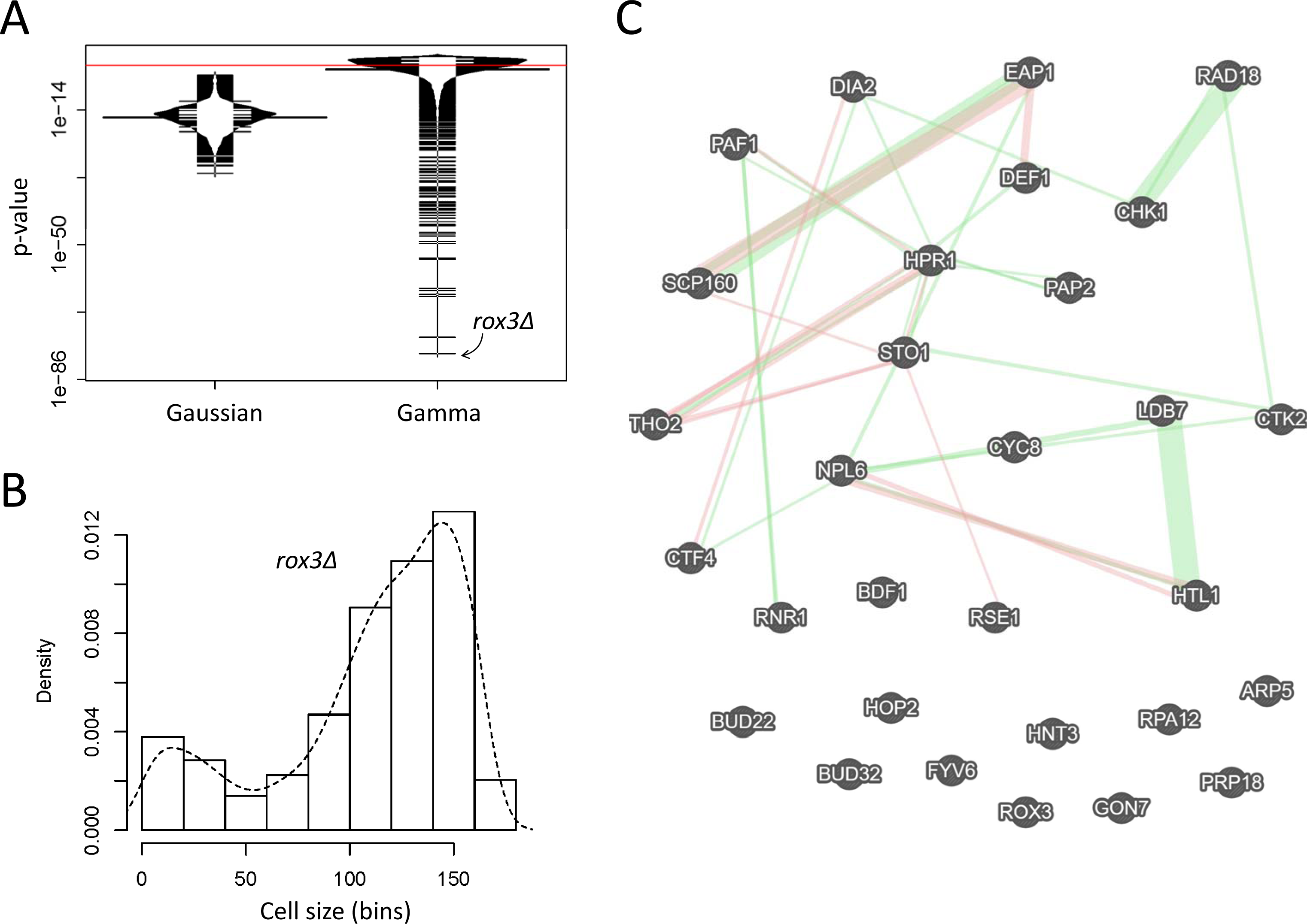
Deletion mut ants of *S. cerevisiae* with severely altered cell size distributions. **A**, Bean plots of the p-values associated with statistical tests of whether cell size values of strains carrying single-gene deletions of all non-essential genes from (Jorgensen et al. 2002) are distributed according to a normal (Gaussian), or Gamma distribution. The statistical tests were performed and displayed as in Figure 2A. The red line indicates a significance level of p=0.01. The most extreme outlier of the Gamma-fitted distributions *(rox3Δ)* is indicated with the arrow. **B**, Histogram and density for the indicated cell size distribution from *rox3Δ* cells, from (Jorgensen et al. 2002). On the y-axis are the density frequency values, while on the x-axis are the cell size bins, encompassing the cell size values shown in the corresponding spreadsheet associated with this plot (see File S1). **C**, Network of the interactions among the 30 genes that belonged in the gene ontology group 'nucleic acid metabolic process' [GO:0090304; p=0.008391). The network was drawn on the GeneMANIA platform (Warde-Farley et al. 2010), with genetic interactions shown in green and physical ones in red.

Next, to identify genes that may be necessary for the gamma distribution pattern of cell size in *S. cerevisiae,* we focused on the samples whose distributions deviated the most from a gamma distribution (i.e., the ones with the lowest p-values shown in Figure 3A, right plot). The sample with the worst fit was from a strain lacking Rox3p, a subunit of the RNA polymerase II Mediator complex (Gustafsson et al. 1997). Not only cells from this mutant were large, as also identified by (Jorgensen et al. 2002), but their size distribution was negatively skewed, with a left-side tail from the main peak (see Figure 3B; the smaller peak to the extreme left of the distribution likely arose from small particulate debris from dead cells in the culture).

Given the severe departure from a gamma distribution for *rox3Δ* cells (Figure 3B), we next looked at the 50 samples with the worst fits. Including *rox3Δ,* these were from 49 deletion strains (one strain in this set was measured twice by (Jorgensen et al. 2002)). The systematic names of these strains are shown in File S1/sheet 'Gamma_deviant_Genes'. Since experimental irregularities could be the reason for the extremely poor fits to a theoretical distribution (e.g., see Figure Sl), we relied on gene ontology enrichment as a functional, unbiased criterion to guide our identification of physiologically relevant mutants. Based on the YeastMine platform (Balakrishnan et al. 2012), 30 of the 49 genes belong to the ontology group 'nucleic acid metabolic process' (GO:0090304; p=0.008391, after Holm-Bonferroni test correction). A smaller group of 16 genes (15 of which were also in the 'nucleic acid metabolic process' set) belonged to the ontology group 'cellular response to DNA damage stimulus' (GO:0006974; p=2.510022e-5 after Holm-Bonferroni test correction). The full gene ontology output for the 30 genes of the 'nucleic acid metabolic process' is shown in File S1/sheet 'GO 0090304'.

Most of the 30 gene products of the 'nucleic acid metabolic process' have a network of previously reported genetic and physical interactions (Figure 3C), consistent with their involvement in common cellular processes. Upon closer inspection, most mutant strains in this group lack genes encoding gene products that regulate gene expression globally, especially transcription *(LDB7, HTL1, NPL6, ROX3, CYC8, PAF1, HPR1, BUD32, CTK2, GON7, RPA12, BDF1, ARP5, THO2),* but also splicing and RNA processing *(RSE1, BUD22, STO1, PRP18, PAP2),* or translation *(SCP160, EAP1).* There was some obvious coherence in this set of gene deletions, in that RSC complex submodule-encoding genes *(LDB7, HTL1, NPL6)* as well as two genes encoding members of the THO complex *(THO2, HPR1)* were identified. To examine if non-gamma cell size distributions were a phenotype common among deletions of each of the components of these large transcription-related complexes, we looked at their corresponding size distributions, for each of the components of the RSC, THO, PAF, and Mediator (MED) complexes interrogated by (Jorgensen et al. 2002). Interestingly, every gene deletion encoding a gene product that is part of the RSC complex had a cell size distribution that deviated significantly from gamma (Figure S3A). In contrast, only a subset of PAF or THO deletions had gamma-deviant size distributions (Figure S3B, C), while most of the MED deletions were similar to wild type, with the notable exception of the extreme distribution of *rox3Δ* cells (Figure S3D).

Next, we looked at the empirical cell size distributions of the corresponding 30 deletion mutants. Every single one had a severely irregular size distribution (Figure 4). Several distributions resembled that of *rox3Δ* cells, with a pronounced negative skew *(scp160Δ, bud32Δ, def1Δ;* Figure 4A), while others were very irregular, even multimodal *(sto1Δ, ctf4Δ, dia2Δ, eap1Δ, arp5Δ, cyc8Δ, fyv6Δ, hpr1Δ, bdf1Δ, npl6Δ, prp18Δ, tho2Δ, htl1Δ, gon7Δ;* Figure 4A). Strikingly, all outliers were also abnormally large cell size mutants (Figure 4A, B). Most had already been identified as such by (Jorgensen et al. 2002) and others (Zhang et al. 2002; Manukyan et al. 2008), but some had not. These previously unidentified large cell size mutants lacked the following genes: *CHK1,* encoding a serine/threonine kinase and DNA damage checkpoint effector (Liu et al. 2000); *RAD18,* encoding an E3 ubiquitin ligase (Bailly et al. 1997); *RPA12,* encoding RNA polymerase I subunit A12.2 (Van Mullem et al. 2002); *RSE1,* encoding a splicing factor (Chen et al. 1998); *STO1,* encoding a large subunit of the nuclear mRNA cap-binding protein complex (Colot et al. 1996); *FYV6,* encoding a protein of unknown function (Wilson 2002); and *PAP2,* encoding a non-canonical poly(A) polymerase (Vanacova et al. 2005).

**FIGURE 4.**
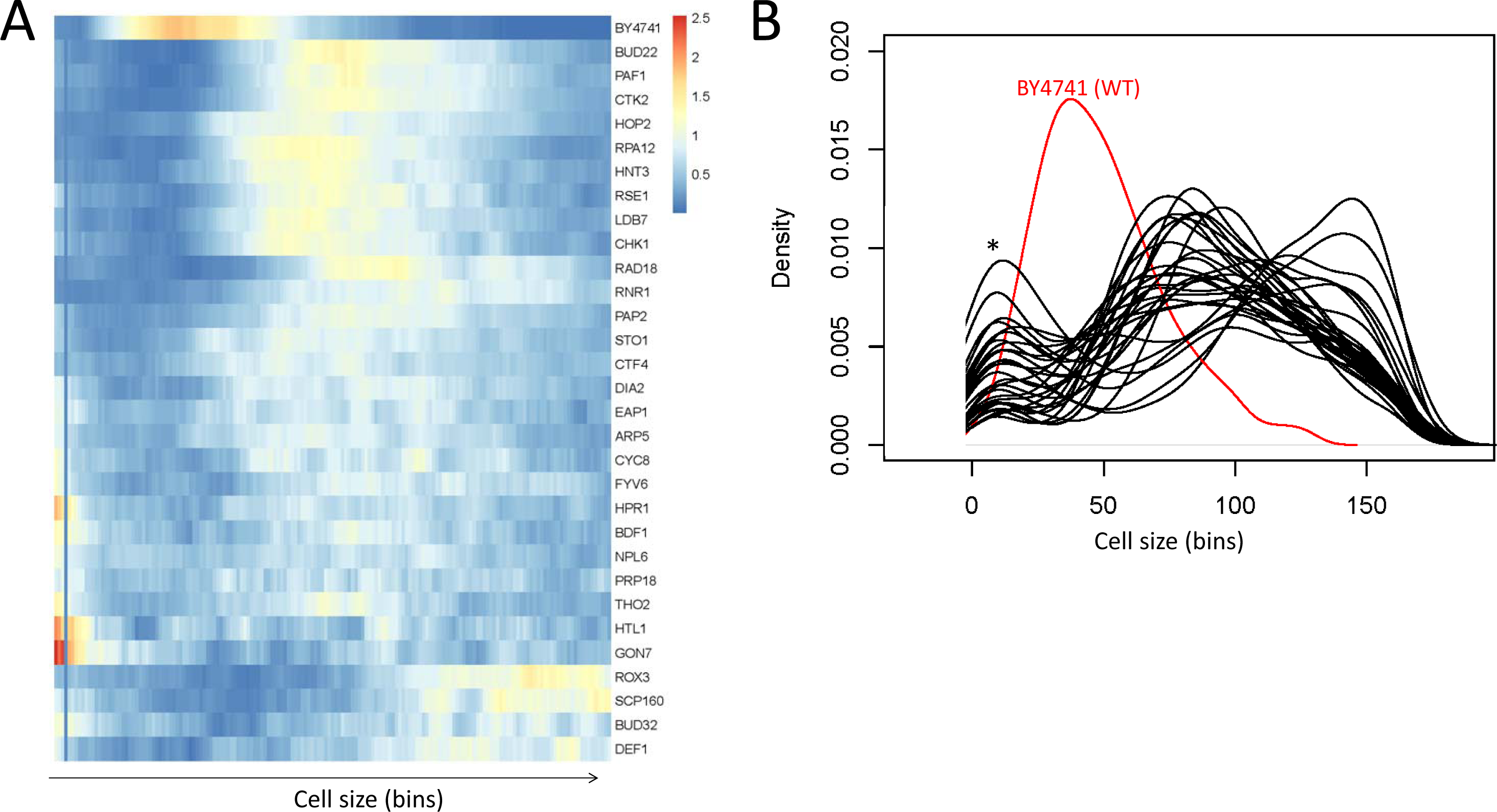
Cell size distributions from the gene deletions that belonged in the gene ontology group 'nucleic acid metabolic process' [GO:0090304]. **A**, Heatmap showing the clustering of cell size density frequencies from the outliers. The frequency values for each mutant are shown along the bins encompassing the cell size values shown in the corresponding spreadsheet associated with this plot (see File S1). The heatmap was generated with the *pheatmap* R language package. **B**, Density plots of cell distributions of the same deletion mutants shown in A. The deletion mutants are shown in black, while their wild type counterpart is shown in red. On the y-axis are the density frequency values, while on the x-axis are the corresponding cell size bins. The asterisk indicates a peak in the distributions that likely arose from small particle debris in the cultures.

To further examine the connection between large cell size and deviation from a gamma distribution, we focused on the deletion strains whose median cell size values were in the top 5% (the criterion used by (Jorgensen et al. 2002), to define their large, *lge,* mutants). Not only were all 49 deletion strains whose distribution differed most significantly from a gamma pattern in this group, but these mutants were also some of the largest ones in the entire collection (Figure S4A, B). Deviations from gamma-distributed cell size values are strongly associated with a very large cell size (p < 2.2E-16; based on the Wilcoxon rank sum test with continuity correction, between the two groups shown in Figure S4A). The remaining mutants with large cell size (shown as 'Other' in Figure S4) were enriched for the gene ontology group “mitotic cell cycle” (GO:0000278; p=1.64E-07), probably reflecting the expected increase in cell size due to a cell cycle block. Hence, it appears that while cell cycle blocks or presumed broad perturbation of gene expression can lead to a larger cell size, it is mostly presumed perturbations to global gene expression or associated processes that lead to deviations from gamma-distributed cell size values.

Next, we looked into the association between poor fitness and deviation from gamma-distributed cell size values. Of the 49 deletion strains whose distribution differed most significantly from a gamma pattern, 31 of the corresponding deletion mutants in a homozygous diploid background had also been reported to have reduced fitness compared to wild type cells in these culture conditions (Giaever et al. 2002). To answer if gamma-deviant mutants were also associated with an extreme reduction in fitness, we compared their fitness scores to the fitness scores of all other remaining 526 mutants with reduced fitness (Figure S4C). The 31 'Gamma_deviant” mutants had an overall significantly poorer fitness than the “Other” 526 mutants ((p = 6.163E-06; based on the Wilcoxon rank sum test with continuity correction). However, the difference in fitness was not as pronounced as was the difference in size (Figure S4C, vs. Figure S4A, respectively). Besides, more than a third (18 out of 49) of the mutants with cell size values that deviated the most from a gamma distribution pattern had a fitness level in these culture conditions that was indistinguishable from wild type, whereas all gamma-deviant mutants were also large size mutants. Hence, we conclude that although deviations from a gamma distribution of cell size values can be associated with poor fitness, the strength of that association is not nearly as great as that with large cell size.

Lastly, we also examined if mutants with extremely small mean cell size *(whi;* the 5% of mutants with the smallest median cell size, as defined by (Jorgensen et al. 2002)) are more likely to deviate from a gamma pattern of cell size distributions. Using the p-values of the Anderson-Darling test as a reference for gamma-deviant distributions, we found that there was no significant difference between *whi* mutants and strains that were not classified as size mutants by (Jorgensen et al. 2002) (p=0.434, based on the Kruskal-Wallis one-way analysis of variance by ranks, followed by the post-hoc Nemenyi test). In contrast, the same analysis looking for deviations from gamma distributions indicated that large cell size mutants *(lge)* are different from strains that were not classified as size mutants by (Jorgensen et al. 2002) (p=0.048, based on the Kruskal-Wallis one-way analysis of variance by ranks, followed by the post-hoc Nemenyi test). While there is a clear association between extreme large size and deviations from gamma distribution we documented (e.g., see Figure S4), the size of mutants that are extremely small do not deviate from a gamma distribution. We illustrate the distribution of one of the most extreme *whi* mutants, *sfp1Δ* cells (Fig. S5), whose distribution fits a gamma distribution (p = 0.211, based on the Anderson-Darling test) as an example (Fig. S5). It is also worth noting that extreme variations in birth size *(sfp1Δ:* 11fL, WT: 22fL; *cln3Δ:* 34fL; calculated as described in (Truong et al. 2013; Soma et al. 2014)) or critical size *(sfp1Δ:* at 73% the size of WT (Jorgensen et al. 2004); *cln3Δ:* at >2-fold the size of WT (Soma et al. 2014)) among these mutants are not necessarily associated with severe deviations from gamma distributions (Figure S5).

### Validation of altered cell size distributions in selected homozygous diploid deletion mutants

In the outlier set of mutants we identified, we were intrigued by the preponderance of gene products connected to transcription. Hence, we decided to validate experimentally the size distributions of strains lacking *PAF1, CTK2, DEF1, BDF1, THO2, PAP2,* or *LDB7.* We used diploid strains carrying homozygous deletions of these genes, to minimize the effects of suppressors that may have been present in the haploid strains used by (Jorgensen et al. 2002). The cell size distributions for all these strains deviated from the gamma distribution of experiment-matched wild type cells (Figure 5; and File S1). These results strengthen the notion that perturbations in the control of gene expression may disrupt the distribution of cell sizes in a population.

**FIGURE 5.**
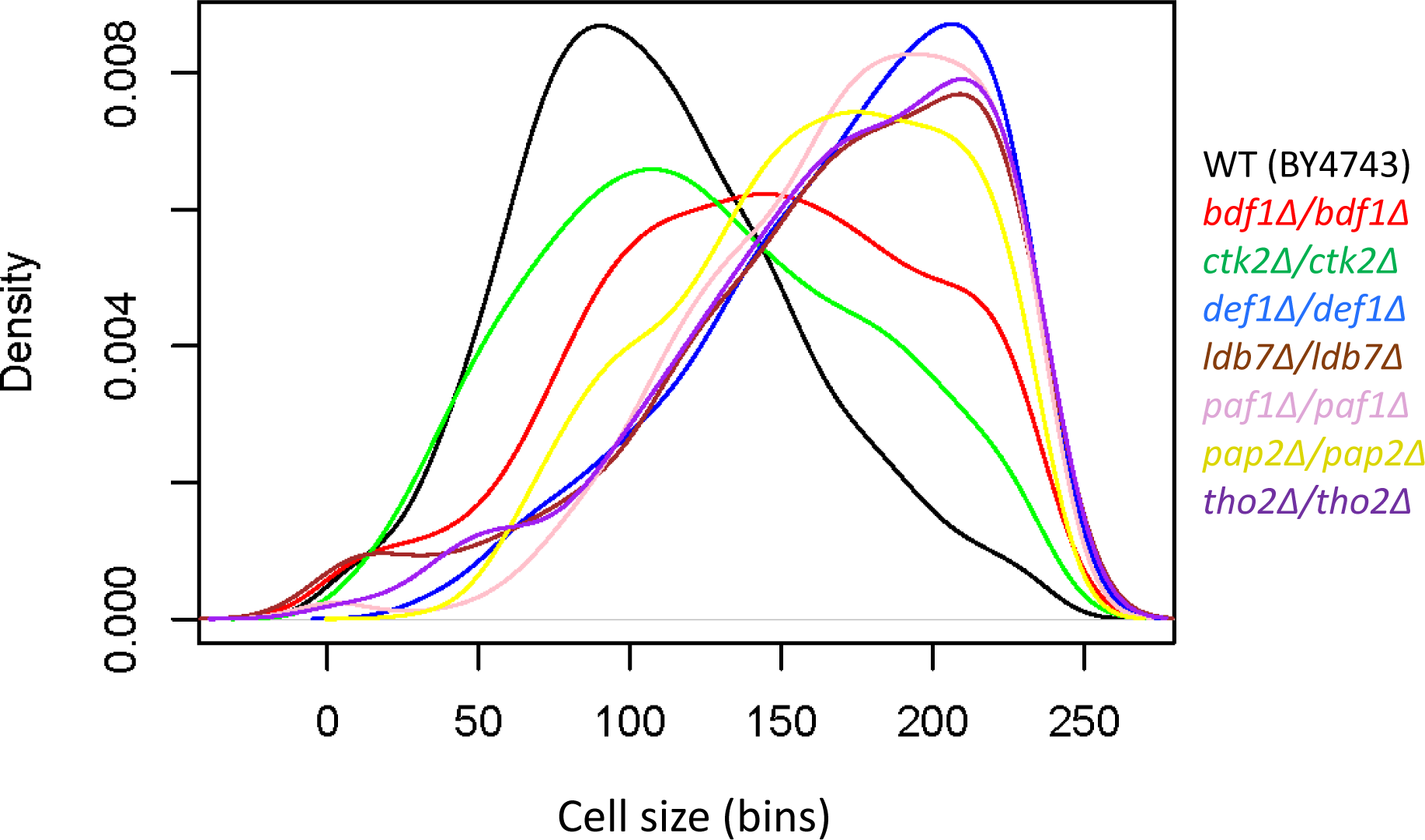
Density plots of cell size distributions from homozygous diploid strains. The cell size of the indicated deletion mutants was measured in standard YPD medium (1% yeast extract, 2% peptone, 2% dextrose), as described in Materials and Methods. Each strain was measured several times (see File S1; sheet 'figure5'), from which representative density plots are shown. On the y-axis are the density frequency values, while on the x-axis are the cell size bins, encompassing the cell size values shown in the corresponding spreadsheet associated with this plot (see File S1).

Lastly, we also measured the DNA content of the strains lacking *PAF1, CTK2, DEF1, BDF1, THO2, PAP2,* or *LDB7,* to ask if their altered cell size distribution is associated with a particular cell cycle profile or ploidy abnormalities. From a genome-wide study, we had previously reported that loss of *PAF1, PAP2* or *LDB7* increased the percentage of cells that are in the G1 phase of the cell cycle (Hoose et al. 2012). Here, we confirmed this phenotype and found that an apparent G1 delay is also the case for cells lacking *CTK2* or *DEF1,* while the loss of *BDF1* or *THO2* does not lead to significant changes in the DNA content (Figure S6A). Hence, it appears that deviations from gamma distribution of cell size values are not obligately associated with a particular cell cycle profile, a conclusion reinforced by additional results we will describe later (see Figure S7).

### Point mutations in the trigger loop of RNA polymerase II alter cell size

It has been proposed that global transcriptional output is tuned with cell size through some poorly characterized mechanism, perhaps by increased RNA Pol II abundance or processivity, or altered mRNA stability in large cells (Zhurinsky et al. 2010; Marguerat and Bahler 2012; Mena et al. 2017; Padovan-Merhar et al. 2015). However, the cell size phenotypes of mutants that affect core RNA polymerase functions are not well-characterized, not least because only four (Rpb4,7,9,12) of the 12 subunits in the complex are non-essential in at least some genetic backgrounds (Myer and Young 1998; Giaever et al. 2002). Cells lacking any one of the non-essential RNA polymerase core subunits have been reported to be large (Jorgensen et al. 2002; Zhang et al. 2002), and usually display a G1 delay (Hoose et al. 2012).

To test the role of global transcription mechanism in cell size control, we examined a set of well-characterized point mutants carrying single amino acid substitutions in the largest Pol II subunit (Rpo21/Rpb1), which are either increased activity (biochemical and genetic “gain of function” GOF, E1103G, G1097D) or decreased activity (biochemical and genetic “loss of function” LOF : H1085Y, N1082S, H1085Q; genetic loss of function: H1085W)(Kaplan et al. 2012; Qiu et al. 2016; Kaplan et al. 2008; Braberg et al. 2013). We found that in all cases cell size was increased, correlating with the extent of catalytic rate alteration (Kaplan et al. 2008; Kaplan et al. 2012) and/or mutant growth rate defects (Kaplan et al. 2012; Malik et al. 2017; Qiu et al. 2016) (Figure 6). Interestingly, albeit the sizes of moderate mutants E1103G, N1082S, and H1085Q were moderately larger than wild type, the distribution pattern did not change (Figure 6). In contrast, the three severe alteration-of-function mutants (G1097D, H1085Y, H1085W) had a very large size, and their distribution deviated from the expected gamma distribution pattern (Figure 6). Pol II LOF and GOF mutants are distinguishable biochemically and genetically, though growth rates of these strains scale with the magnitudes of their biochemical defects and extent of their gene expression defects and genetic interactions. Similarly, Pol II mutant cell sizes, regardless of LOF or GOF status, correlate with their growth rates. We conclude that altering global transcription, severe or moderate, and in either direction, with gain-or loss-of-function Pol II mutations, increases cell size. Furthermore, severe alteration-of-function Pol II mutations abrogate the gamma distribution of cell size values.

**FIGURE 6.**
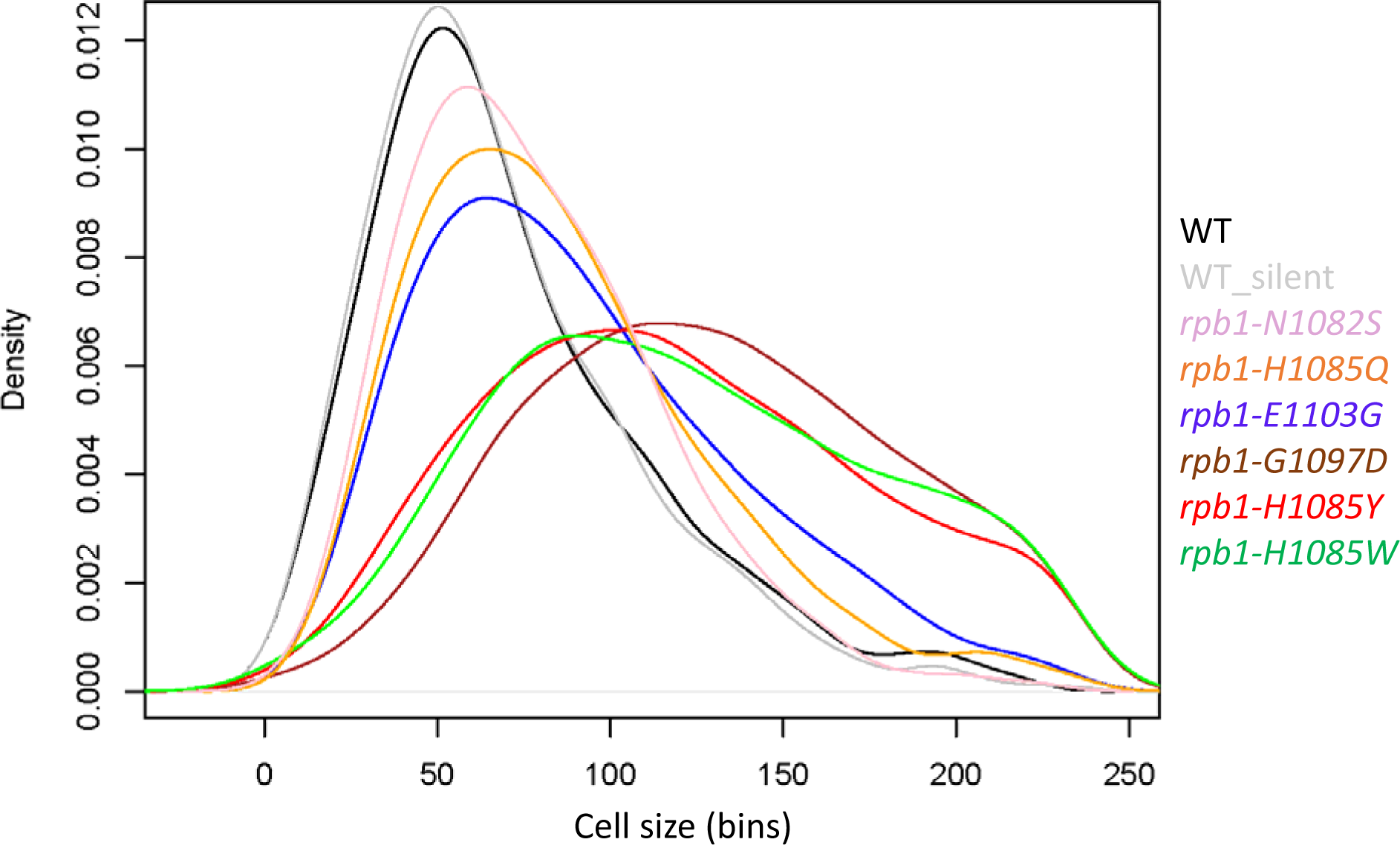
Density plots of cell size distributions from Pol II trigger-loop point mutants. The cell size of the indicated mutants was measured in standard YPD medium (1% yeast extract, 2% peptone, 2% dextrose), as described in Materials and Methods. Each strain was measured several times (see File S1; sheet 'figure6'), from which representative density plots are shown. On the y-axis are the density frequency values, while on the x-axis are the cell size bins, encompassing the cell size values shown in the corresponding spreadsheet associated with this plot (see File S1).

Next, we measured the DNA content of these Pol II mutants. Similarly to the deletion mutants that we analyzed in Figure S6A, the severe alteration-of-function *rpb1* mutants (G1097D, H1085Y, H1085W) displayed a significant increase in the G1 DNA content (Figure S6B). Interestingly, both substitutions at position 1085 (Y or W) also displayed a cell cycle profile consistent with S-phase delay, because the peaks corresponding to un-replicated and replicated DNA were not separated (Figure S6B). The two moderate loss-of-function mutants (N1082S, H1085Q) had a modest increase of cells with G1 DNA content. The gain-of-function mutant (E1103G) did not display G1 or S-phase delay (Figure S6B). If anything, there was a slight increase of the G2/M DNA content in *rpb1-E1103G* cells, which along with their slightly larger cell size (Figure 6) and moderately slower proliferation rates (Kaplan et al. 2012; Malik et al. 2017), argues for a possible mitotic delay in this mutant. A potential mitotic delay could be consistent with Pol II increased activity mutants showing increased rates of minichromosome loss in chromosome segregation experiments (Braberg et al. 2013).

### Altered cell cycle progression is not sufficient to alter the gamma distribution of cell size values

The altered cell cycle profiles of Pol II mutants and strains lacking genes involved in transcription (Figure S6) raised the question of whether abnormalities in cell cycle progression are the cause of the poor fits of cell size values to a gamma distribution in many of these mutants. To test this possibility, we examined the goodness-of-fit to a gamma distribution, using the Anderson-Darling associated p-values, for the following groups of deletion strains (Figure S7): Mutants displaying a 'High G1' DNA content, usually associated with a G1 delay (Hoose et al. 2012); mutants displaying a 'Low G1' DNA content, usually associated with a G2/M delay (Hoose et al. 2012); and mutants lacking genes of the DNA damage checkpoint biological process (Gene ontology group G0:0000077). These groups were compared to each other and all remaining strains analyzed by (Jorgensen et al. 2002). While some outliers in these groups had altered cell size distributions *(e.g.,* cells lacking the Chk1p checkpoint kinase in the G0:0000077 group), there was no statistically significant difference among these groups of mutants (based on the non-parametric Kruskal-Wallis test, p=0.4841). Hence, cell cycle defects observed in several transcription mutants are not sufficient for explaining the significant deviations from the gamma distribution of cell size values in these strains. Instead, it is likely that a constellation of defects in gene expression or defects linked to transcriptional impact to the genome is the cause of cell size distribution derangement.

## DISCUSSION

We discuss our results that cell size values in *S. cerevisiae* follow a gamma distribution and the role of global transcription in the control of cell size.

In biology, lognormal and gamma distributions have been proposed to describe tissue growth models (Mosimann 1988). In both cases, the observed distributions are thought to arise from random fluctuations of many independent variables. Lognormal patterns reflect an aggregate multiplicative process generated from exponential patterns of growth (Koch 1966; Frank 2009). Despite random fluctuations, the growth of the overwhelming majority of cellular components is influenced proportionally, leading to log-normality (Koch 1966; Koch and Schaechter 1962). Similarly, gamma distributions represent the aggregate of many power-law and exponential processes (Frank 2009). With regards to cell size control, it is important to note that not only wild type cells but also size mutants, large or small, appear to maintain their size in a given environment. Such stationary size distributions, with their narrow range of coefficients of variation across populations with different mean size (Anderson et al. 1969), are accommodated by the properties of lognormal and gamma distributions (Hosoda et al. 2011; Dennis and Patil 1984; Kilian et al. 2005; Frank 2016).

Additionally, our data support the notion that global transcription mechanisms are necessary to balance expression with cell size (Figures 3-6), as postulated previously (Zhurinsky et al. 2010; Marguerat and Bahler 2012; Mena et al. 2017; Padovan-Merhar et al. 2015). The relationships between cell size and mRNA synthesis and decay rates are complex. It has been proposed for budding yeast that alterations to synthesis can correspondingly be buffered by changes in mRNA decay, and vice versa, enabling cells to maintain gene dosage in the face of global perturbations to gene expression (Sun et al. 2013; Sun et al. 2012; Haimovich et al. 2013). Cell size and genome replication are sources of potential perturbations to gene expression because an increase in cell volume will dilute concentrations of cellular factors unless compensated by global changes. Conversely, during replication, the dosage of the genome per cell doubles which can be countered by a concomitant increase in cell size. Prior work indicated that a subset of factors involved in mRNA turnover could also generate large cells. Here, we show that a number of additional mutants in genes with potential widespread roles in gene expression, including alterations to the Pol II active site, lead to larger cells that can have altered distributions of sizes compared to wild type. A question that arises from this work is whether perturbed gene expression deregulates specific factors that control cell size homeostasis, or an increase in cell size is a consequence of globally defective gene expression, whereby alterations to global expression processes elicit buffering mechanisms that function through changes in cell volume. While an extreme size deviation is common among the gamma distribution-deviating strains, the actual distributions of these mutants appear to be of more than one class, suggesting either complex or distinct underlying mechanisms.

How could perturbations in global transcription alter the observed gamma distribution of cell size? In live cells, it appears that constitutive gene expression occurs stochastically in bursts at the single molecule level, with the burst magnitude and frequency leading to gamma-distributed, steady-state levels of the produced protein (Cai et al. 2006; Li and Xie 2011; Friedman et al. 2006). Changes in the burst magnitude and frequency of transcription events have been proposed to explain the scaling of mRNA counts with cell size (Padovan-Merhar et al. 2015). The reproductive property of the gamma distribution predicts that if the independent random variables themselves are gamma-distributed, then the aggregate of all these random variables will also be gamma-distributed (Johnson 1994). Cell size is routinely viewed as a proxy for cell mass (Turner et al. 2012). Cell mass, in turn, is mostly determined by the accumulated macromolecules, especially proteins (Lange and Heijnen 2001). Hence, it is reasonable to speculate that global perturbations in the mechanics of gene expression that cause the steady-state levels of individual gene products to deviate from their gamma-distributed pattern could also perturb the aggregate gamma pattern of gene expression and its manifestation in cell size. Perturbation of global transcription through altered Pol II catalytic activity leads to changes in cell size, consistent with such a model. Conversely, a subset of specific gene expression perturbations may in aggregate lead to altered cell physiology that results in both extreme cell size and deviation from a gamma distribution.

We show here that some mutants are not just extremely large, their populations show distribution changes from wild type. Such changes in the distribution of sizes may occur for any number of reasons. Factors observed here may elicit cell size alterations directly through gene expression changes or indirectly through mRNA export defects or transcription-dependent DNA damage or recombination (TREX/THO complex members *hpr1Δ, tho2Δ* (Prado et al. 1997; Piruat and Aguilera 1998)), reduced ability to degrade stalled Pol II *(def1Δ* (Woudstra et al. 2002)), or widespread changes to transcription-dependent chromatin modifications or elongation control *(paf1Δ* (Van Oss et al. 2017)) or chromatin structure (RSC submodule components *1db7Δ, htl1Δ, npl6Δ* (Wilson et al. 2006; Cairns et al. 1996)). Our DNA content data (see Figure S6) are not consistent with extreme chromosomal rearrangements in the mutants we examined. We also found that both haploid and diploid size distributions appear to fit better a gamma pattern (Table 1). However, we cannot exclude the possibility that possible aneuploidy or spontaneous diploidization in some of these mutants (especially RSC mutants (Sing et al. 2018), and *def1Δ* cells (Stepchenkova et al. 2018)) may contribute to deviations from gamma distributions. For Pol II mutants, sizes of cells correlate with several phenotypes: strain growth rates, biochemical and genetic defects, and the extent to which a specific mRNA's half-life was increased (Malik et al. 2017), raising intriguing questions about potential causative relationships between these phenotypes. Future experiments monitoring transcription events in single cells of different size could shed light in the relationship between global gene expression and size control.

## Author Contributions

Methodology, C.D.K., M.P.; Formal Analysis, M.P.; Investigation, N.M. H.M.B., J.A., C.D.K., M.P.; Resources, C.D.K., M.P.; Data Curation, M.P.; Writing – Original Draft, M.P., Writing – Review and Editing, C.D.K., M.P.; Visualization, M.P.; Supervision, C.D.K., M.P., Funding Acquisition, C.D.K., M.P.

We thank Marta Karas (Johns Hopkins University) for helpful suggestions for the R code we used. This work was supported by grants from the National Institutes of Health to M.P. (R01GM123139) and C.D.K. (R01GM097260), and the Welch Foundation to C.D.K. (A-1763).

---

**FIGURE S1.**
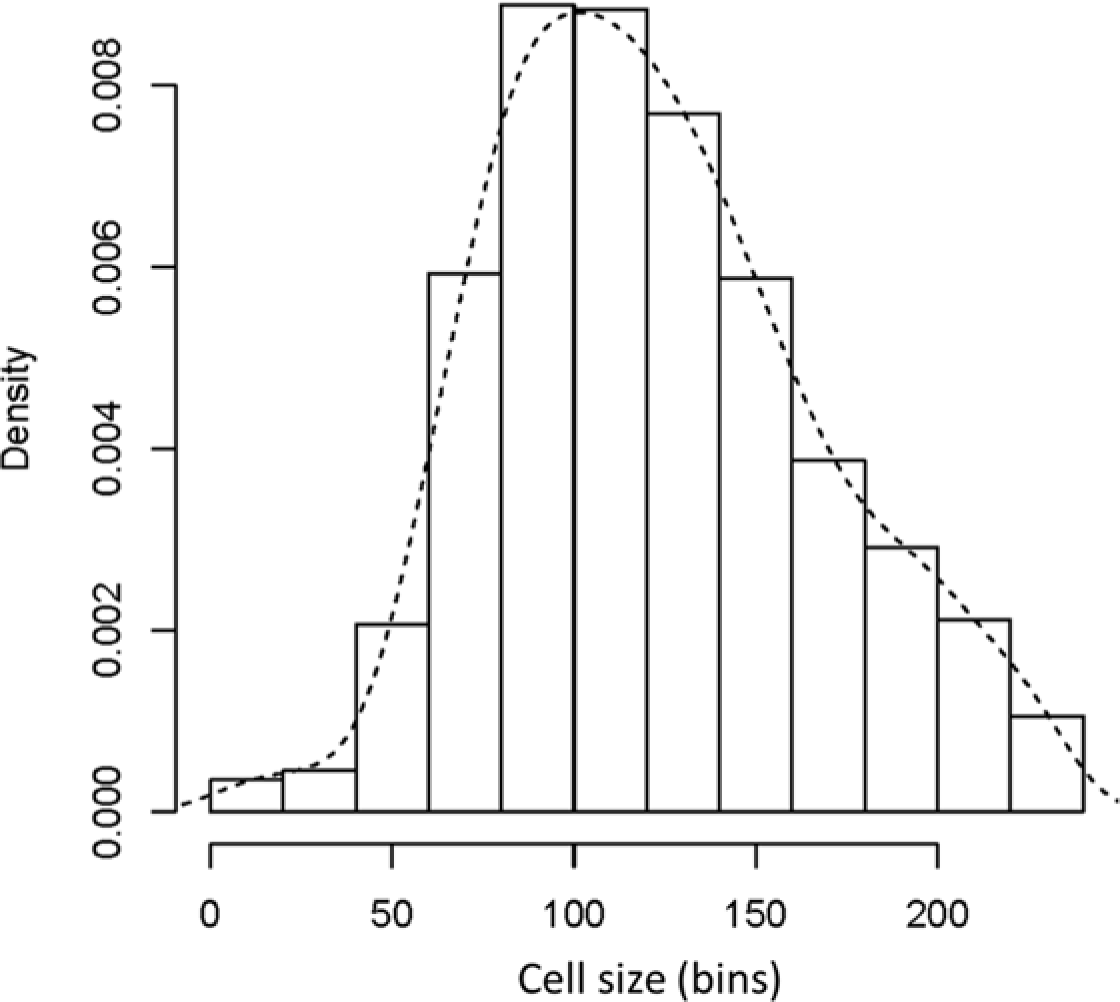
An empirical density distribution of cell size values from wild type cells that did not fit a Gamma distribution. **A**, Histogram and density for the indicated cell size distribution from wild type diploid (BY4743 strain) cells, cultured in standard YPD medium (1% yeast extract, 2% peptone, 2% dextrose). This distribution was from the sole outlier of wild type cells that did not fit a Gamma distribution, from Figure 2A, due to a right-side 'shoulder' in the density. On the y-axis are the density frequency values, while on the x-axis are the cell size bins, encompassing the cell size values shown in the corresponding spreadsheet associated with this plot (see File S1).

**FIGURE S2.**
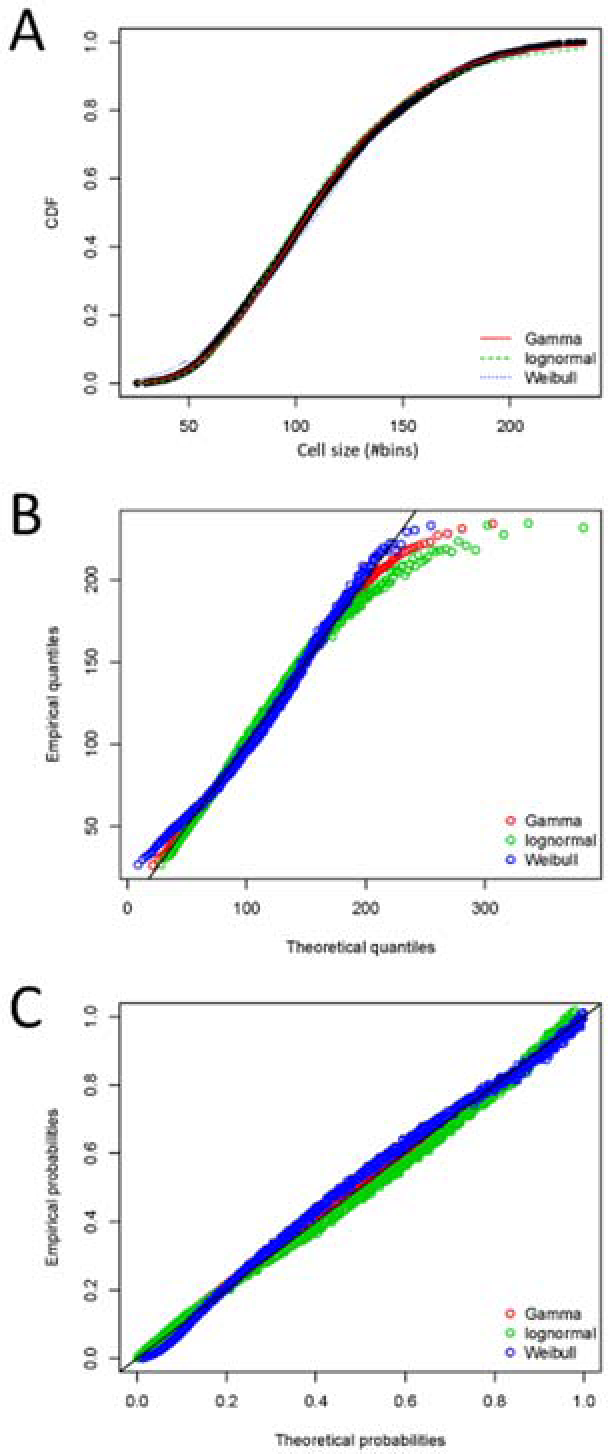
Goodness-of-fit plots for the indicated distributions. **A**, From the same dataset of wild type BY4743 cells shown in Fig. 2, the cumulative distribution function (CDF) of experimental cell size values (shown black), along with the theoretical CDFs of the Gamma, lognormal, and Weibull distributions (shown in red, green and blue, respectively). **B**, Quantile-Quantile (Q-Q) plots, comparing the experimental quantiles of cell size values against the theoretical quantiles of the Gamma, lognormal, and Weibull distributions. Q-Q plots emphasize poor fit at the tails of the distribution. **C**, Probability-Probability (P-P) plots of the CDF of experimental cell size values plotted against the theoretical CDFs of the Gamma, lognormal, and Weibull distributions. P-P plots emphasize poor fit at the center of the distribution.

**FIGURE S3.**
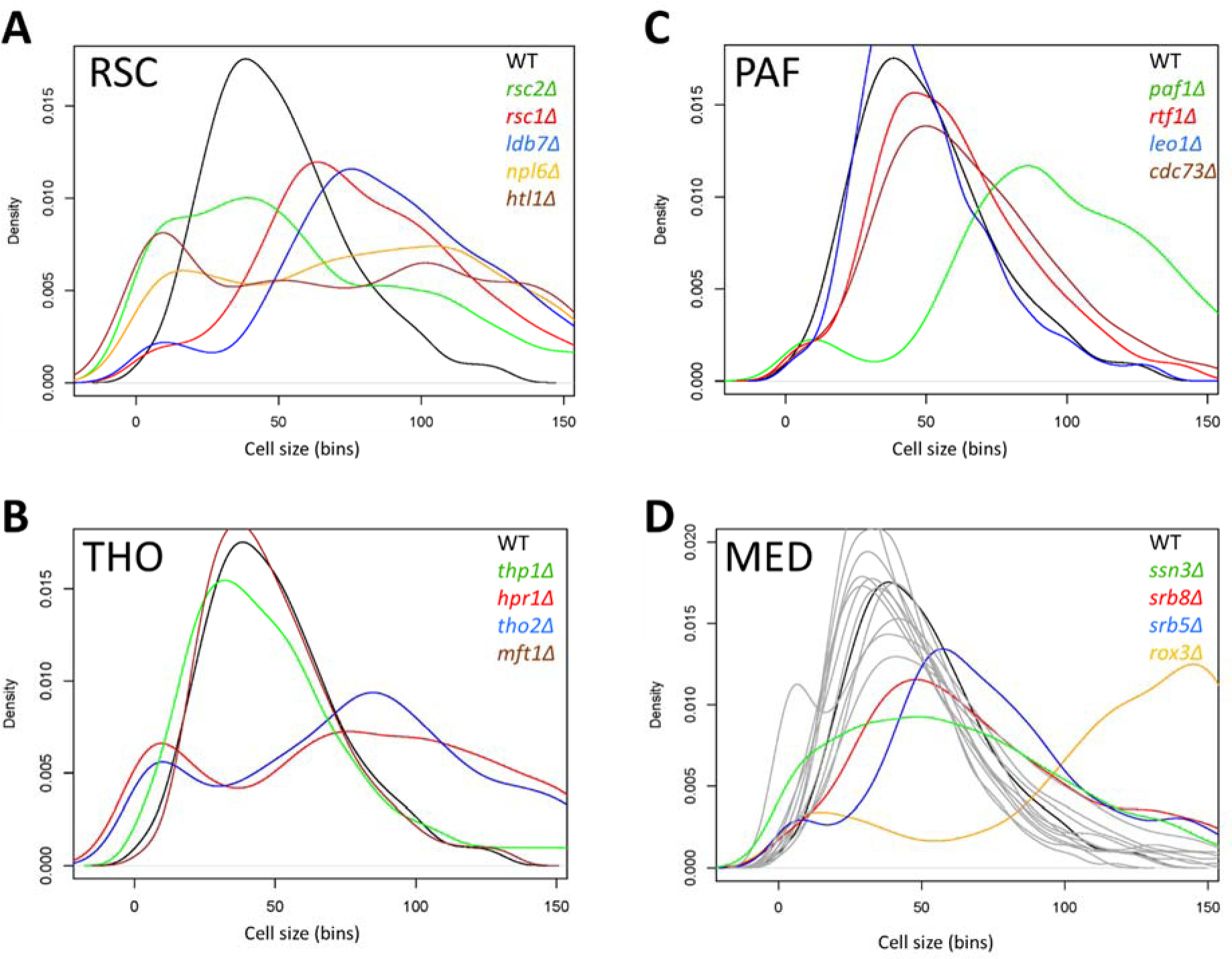
Density plots of cell size distributions from deletions of components of the indicated transcription-related complexes. **(A-D)**. The deletion mutants of components from each complex are the viable ones queried by (Jorgensen et al. 2002). On the y-axis are the density frequency values, while on the x-axis are the cell size bins, encompassing the cell size values shown in the corresponding spreadsheet associated with this plot (see File S1). The wild type is shown in black in every panel. The density plots shown in grey in (D), are from deletions of genes encoding components of the Mediator (MED) complex that had distributions very similar to wild type, and they corresponded to the following genes: *MED1, MED2, PGD1, NUT1, CSE2, SSN2, GAL11, SIN4, SRB2, SOH1, SSN8.*

**FIGURE S4.**
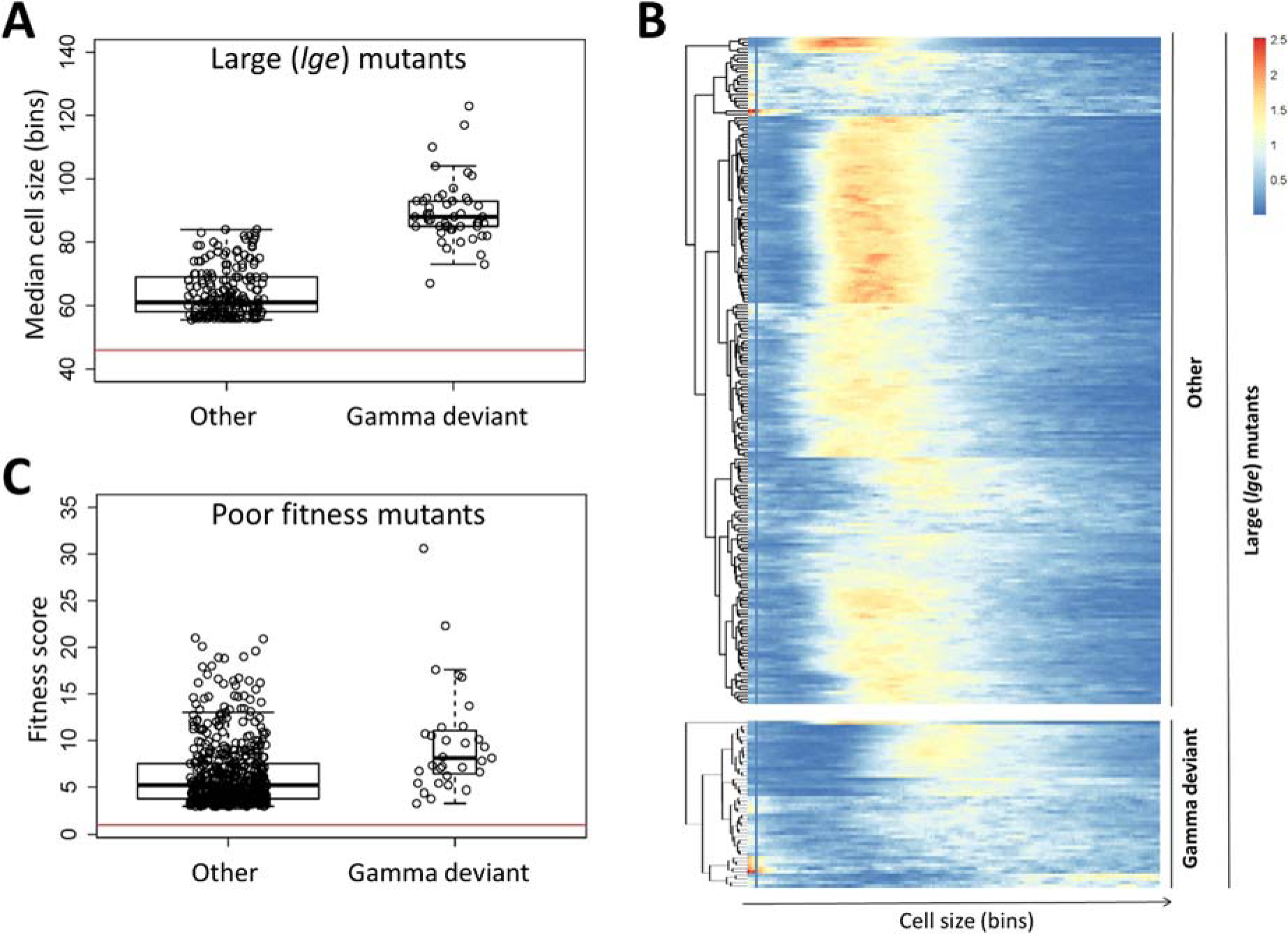
Association of poor fit to gamma distribution of cell size values with large cell size (A, B) or reduced growth fitness (C). **A**, Only a subset of large cell size mutants *(lge)* have distributions that deviate from a gamma pattern. The 253 deletion mutants analyzed by (Jorgensen et al. 2002), which had a median cell volume in the top 5% of all strains that were examined, were grouped in two groups: In the 'Gamma deviant' group were the 49 deletion mutants whose distribution deviated the most from gamma (see File S1), and they were all in the large size mutant group, with many of them being the largest ones. The 'Other' group had the remaining large size mutants. On the y-axis is the median cell size values (in bins) for each of these strains in both groups, while the red line indicates the median value for the wild type samples. The difference in size between the two groups, 'Other' vs. 'Gamma deviant', was highly significant (p < 2.2E-16; based on the Wilcoxon rank sum test with continuity correction). **B**, Heatmap showing the unsupervised hierarchical clustering of cell size density frequencies from the same mutants as in A. The heatmap was generated and displayed as in Figure 4. **C**, The 557 homozygous diploid deletion mutants identified by (Giaever et al. 2002), which had scores indicative of reduced growth fitness, were grouped in two groups: In the 'Gamma deviant' group were the corresponding 31 haploid deletion mutants, which were among the 557 gene deletions identified by (Giaever et al. 2002) to be associated with reduced fitness, and also had size distributions that deviated the most from gamma (see File S1). The 'Other' group had the remaining 526 poor fitness mutants. On the y-axis is the fitness score values (as defined by (Giaever et al. 2002)) for each of these strains in both groups, while the red line indicates the fitness score for the wild type. The difference in fitness between the two groups, 'Other' vs. 'Gamma deviant', was significant (p = 6.163E-06; based on the Wilcoxon rank sum test with continuity correction).

**FIGURE S5.**
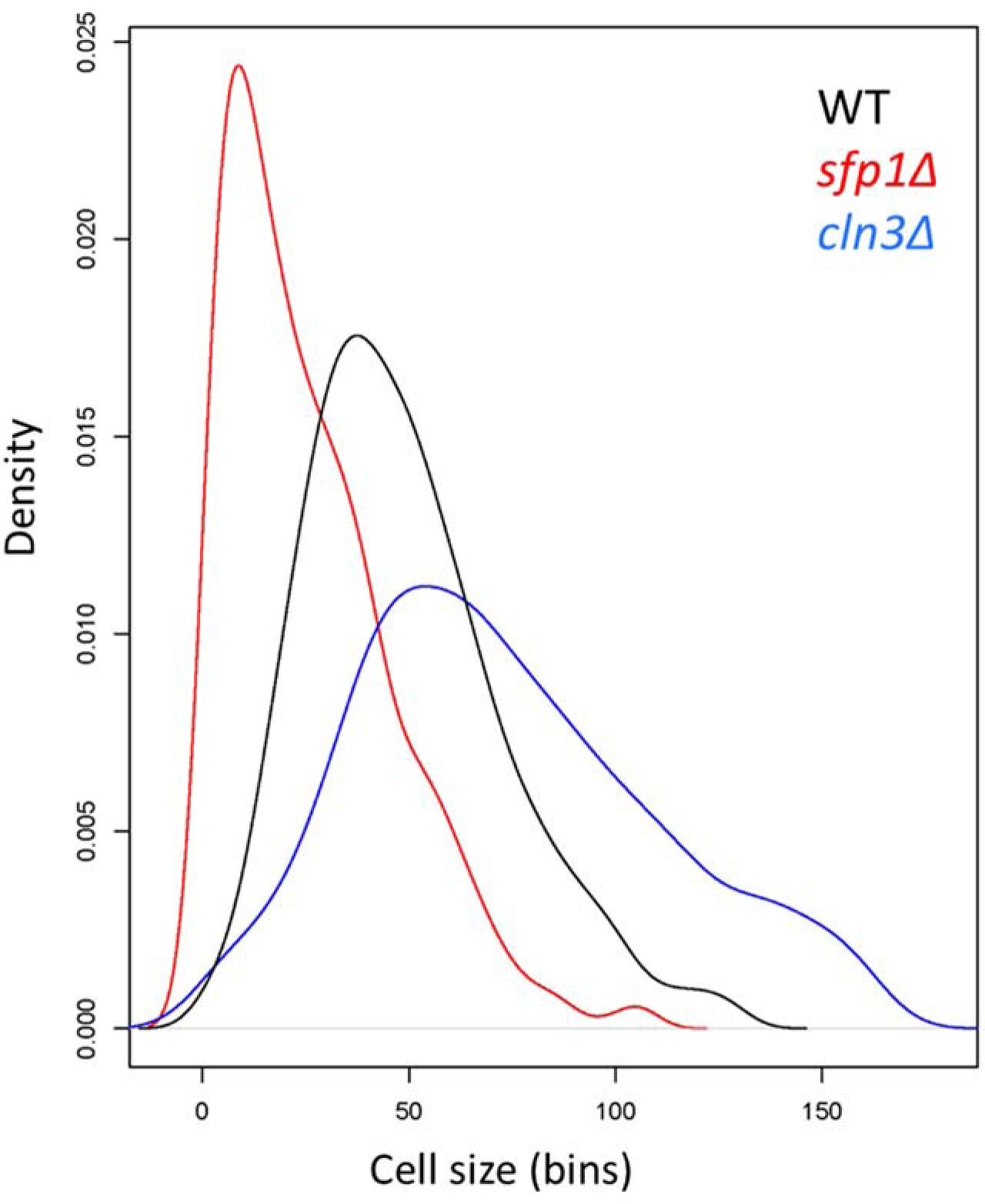
Density plots of cell size distributions from *sfp1Δ* and *cln3Δ* cells. The deletion mutants of a prototypical *whi (sfp1Δ)* and *Ige (c1n3Δ)* were analyzed based on the data from (Jorgensen et al. 2002). On the y-axis are the density frequency values, while on the x-axis are the cell size bins, encompassing the cell size values shown in the corresponding spreadsheet associated with this plot (see Supplementary File 1).

**FIGURE S6.**
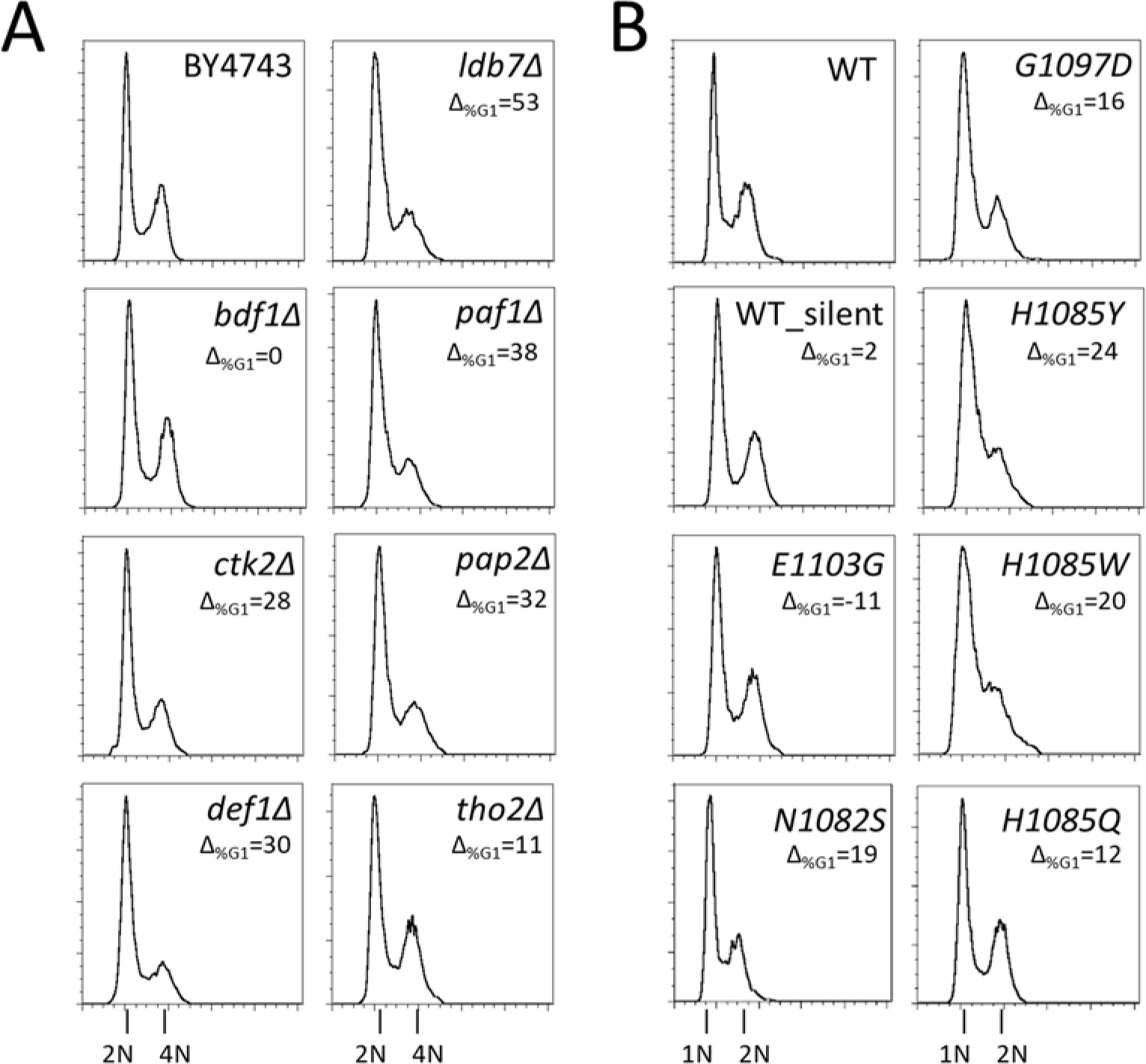
DNA content analysis. In **A**, and **B**, are the DNA content histograms from the same cultures of the strains shown in Figures 5, and 6, respectively. On the x-axis of each panel is fluorescence per cell, and on the y-axis the numbers of cells. The difference in the percentage of cell in the G1 phase of the cell cycle (Δ_%oG1_) compared to experiment-matched wild type cells is shown for each mutant strain, and it is the average of at least two independent measurements.

**FIGURE S7.**
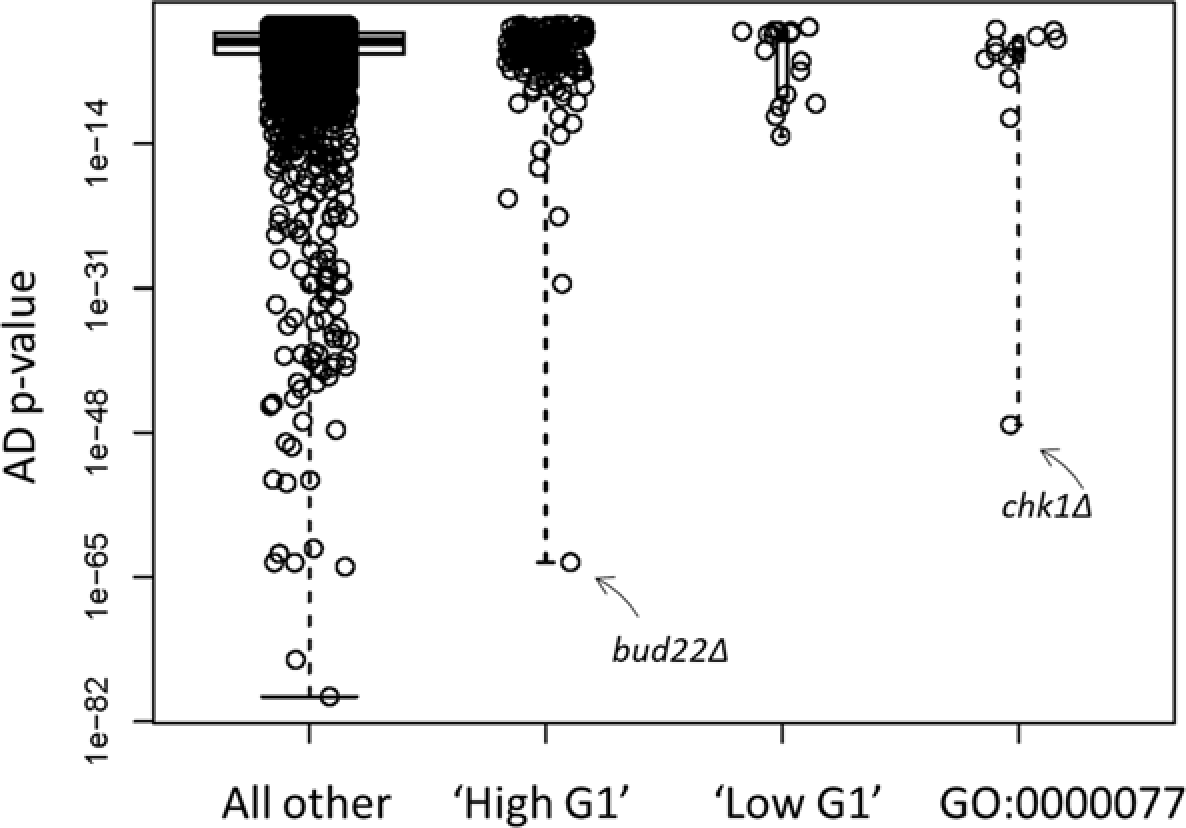
Cells with 'High G1', 'Low G1' DNA content, or carrying deletions of genes in the gene ontology process GO:0000077 (DNA damage checkpoint) have cell size profiles with similar Anderson-Darling associated p-values for gamma distribution fits. The deletion strains with 'High' or 'Low' G1 DNA content were those identified in (Hoose et al. 2012). The G0:0000077 group contained the cell size profiles of strains lacking any single non-essential gene belonging to this ontology group and analyzed by Jorgensen et al. The 'All other” group were all other mutants analyzed by (Jorgensen et al. 2002), which were not in the groups of mutants shown in the plot. The cell size distributions of the corresponding deletion mutants in each case were analyzed as in Figure 2A, fitted to a gamma distribution, with the p-values from the Anderson-Darling test (AD p-value) representing each data point. The groups were not statistically different from each other (Kruskal-Wallis rank sum test, p=0.4841).

**Table S1.**
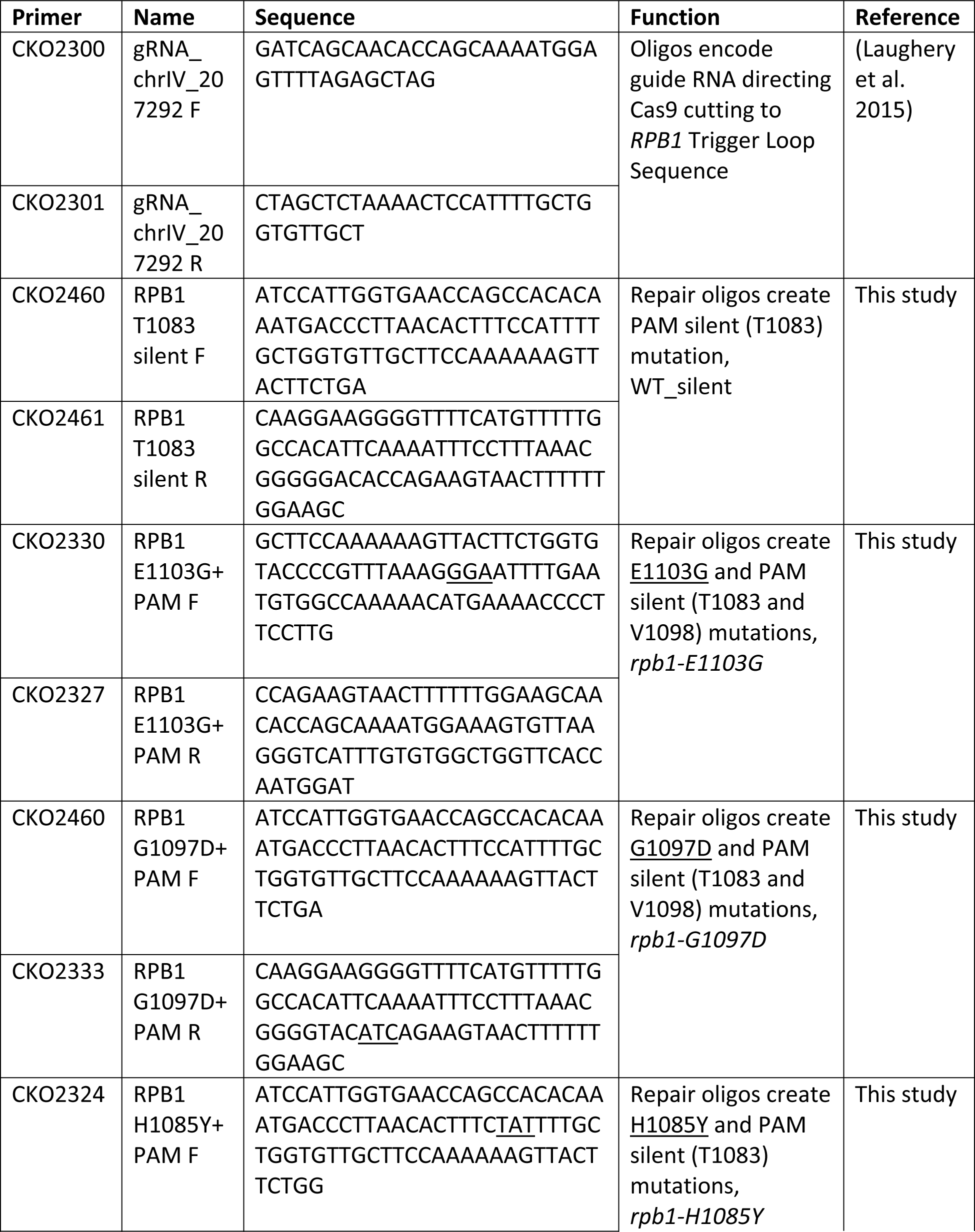
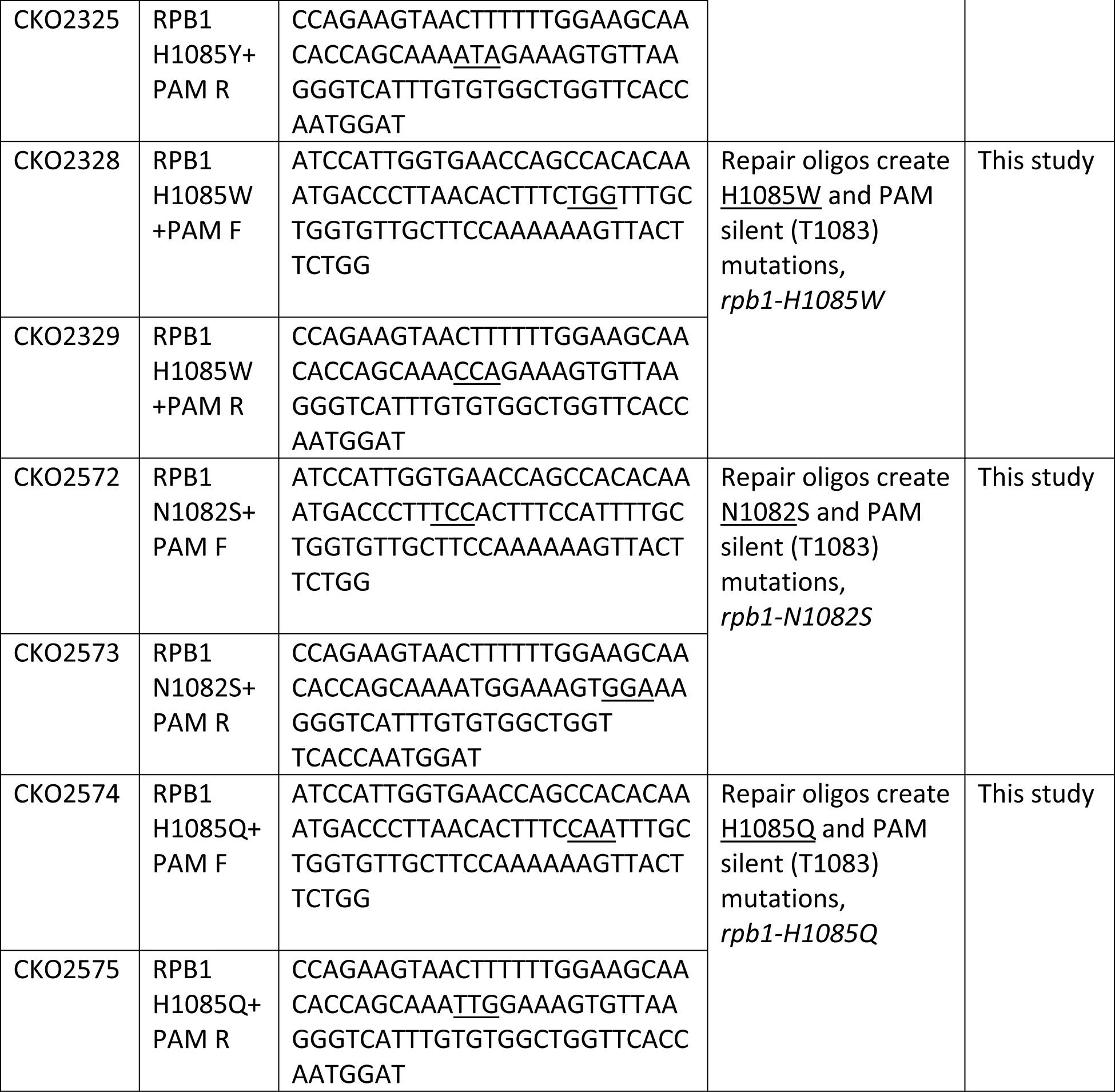
List of oligonucleotides used in this study

**Table S2.** List of yeast strains used in this study.

| **Yeast strain** | **Relevant mutation** | **Genotype** | **Reference** | **Relevant Figure** |
| --- | --- | --- | --- | --- |
| CKY3284 | *RPB1* WT | *MATα ura3-52 his4Δ200 leu2Δ1 or Δ0 trp1Δ63 met15Δ0 lys2-128δ gal10Δ56 RPB1::3XFLAG::kanMX* | This study | Figure 6 and S5 |
| CKY3621 | *RPB1* WT_silent (T1083 silent) | *MATα ura3-52 his4Δ200 leu2Δ1 or Δ0 trp1Δ63 met15Δ0 lys2-128δ gal10Δ56 RPB1_T1083silent::3XFLAG::kanMX* | This study | Figure 6 and S5 |
| CKY3628 | *rpb1E1103G* (T1083 & V1098 silent) | *MATα ura3-52 his4Δ200 leu2Δ1 or Δ0 trp1Δ63 met15Δ0 lys2-128δ gal10Δ56 rpb1-E1103G::3XFLAG::kanMX* | This study | Figure 6 and S5 |
| CKY3624 | *rpb1G1097D* (T1083 & V1098 silent) | *MATα ura3-52 his4Δ200 leu2Δ1 or Δ0 trp1Δ63 met15Δ0 lys2-128δ gal10Δ56 rpb1-G1097D::3XFLAG::kanMX* | This study | Figure 6 and S5 |
| CKY3425 | *rpb1H1085Y* (T1083 silent) | *MATα ura3-52 his4Δ200 leu2Δ1 or Δ0 trp1Δ63 met15Δ0 lys2-128δ gal10Δ56 rpb1-H1085Y::3XFLAG::kanMX* | This study | Figure 6 and S5 |
| CKY3427 | *rpb1H1085W* (T1083 silent) | *MATα ura3-52 his4Δ200 leu2Δ1 or Δ0 trp1Δ63 met15Δ0 lys2-128δ gal10Δ56 rpb1-H1085W::3XFLAG::kanMX* | This study | Figure 6 and S5 |
| CKY3859 | *rpb1N1082S* (T1083 silent) | *MATα ura3-52 his4Δ200 leu2Δ1 or Δ0 trp1Δ63 met15Δ0 lys2-128δ gal10Δ56 rpb1-N1082S::3XFLAG::kanMX* | This study | Figure 6 and S5 |
| CKY3861 | *rpb1H1085Q* (T1083 silent) | *MATα ura3-52 his4Δ200 leu2Δ1 or Δ0 trp1Δ63 met15Δ0 lys2-128δ gal10Δ56 rpb1-H1085Q::3XFLAG::kanMX* | This study | Figure 6 and S5 |

**Table S3.** List of plasmids used in this study.

| Plasmid | Description | Genotype | Reference |
| --- | --- | --- | --- |
| pML107 | Vector contains Cas9 gene and guide RNA Expression cassette | *LEU2* 2μ ori amp<sup>r</sup> Cas9 | (Laughery et al. 2015) |
| pCK2411 | gRNA_<br>chrIV_207292<br>cloned in<br>pML107 | *LEU2* 2μ ori amp<sup>r</sup> Cas9 gRNA_<br>chrIV_207292 | This study |

